# Human and mouse essentiality screens as a resource for disease gene discovery

**DOI:** 10.1101/678250

**Authors:** Pilar Cacheiro, Violeta Muñoz-Fuentes, Stephen A. Murray, Mary E. Dickinson, Maja Bucan, Lauryl M.J. Nutter, Kevin A. Peterson, Hamed Haselimashhadi, Ann M. Flenniken, Hugh Morgan, Henrik Westerberg, Tomasz Konopka, Chih-Wei Hsu, Audrey Christiansen, Denise G. Lanza, Arthur L. Beaudet, Jason D. Heaney, Helmut Fuchs, Valerie Gailus-Durner, Tania Sorg, Jan Prochazka, Vendula Novosadova, Christopher J. Lelliott, Hannah Wardle-Jones, Sara Wells, Lydia Teboul, Heather Cater, Michelle Stewart, Tertius Hough, Wolfgang Wurst, Radislav Sedlacek, David J. Adams, John R. Seavitt, Glauco Tocchini-Valentini, Fabio Mammano, Robert E. Braun, Colin McKerlie, Yann Herault, Martin Hrabě de Angelis, Ann-Marie Mallon, K.C. Kent Lloyd, Steve D.M. Brown, Helen Parkinson, Terrence F. Meehan, Damian Smedley, on behalf of the Genomics England Research Consortium and the International Mouse Phenotyping Consortium

## Abstract

Although genomic sequencing has been transformative in the study of rare genetic diseases, identifying causal variants remains a considerable challenge that can be addressed in part by new gene-specific knowledge. Here, we integrate measures of how essential a gene is to supporting life, as inferred from the comprehensive viability and phenotyping screens performed on knockout mice by the International Mouse Phenotyping Consortium and from human cell line essentiality screens. We propose a novel, cross-species gene classification across the <u>Fu</u>ll <u>S</u>pectrum of Intolerance to <u>L</u>oss-of-function (FUSIL) and demonstrate that genes in five mutually exclusive FUSIL categories have differing characteristics in the biological processes they regulate, tissue expression levels and human mutation rates. Most notably, Mendelian disease genes, particularly those associated with developmental disorders, are highly overrepresented in the developmental lethal category, representing genes not essential for cell survival but required for organism development. Exploiting this finding, we have screened developmental disorder cases from three independent disease sequencing consortia and identified potentially pathogenic, *de novo* variants shared in different patients for several developmental lethal genes that have not previously been associated with rare disease. We therefore propose FUSIL as an efficient resource for disease gene discovery.

## INTRODUCTION

Discovery of the genetic causes of monogenic disorders has shifted from genetic analysis of large cohorts or families with the same phenotype, to a genotype-driven approach that is able to identify ultra-rare variants associated with a disorder in one or few individuals. Published studies by the Centers for Mendelian Genomics^1^, Deciphering Developmental Disorders^2, 3^ and the Undiagnosed Disease Network^4^ have successfully used whole-exome or genome sequencing to find the causal variant in up to 40% of patients. However, the majority of cases remain unsolved and metrics incorporating gene-level information can help to identify candidate variants in previously unknown disease genes that are subsequently confirmed as causative in functional *in vitro* and *in vivo* studies. Measures of genetic intolerance to functional variation, based on whole exome and genome sequencing data from broad populations of healthy individuals or cohorts affected by non-severe and non-pediatric disease, represent one class of metrics that have been used to prioritise candidate disease genes where heterozygous, dominant effects are suspected^5–7^. Use of standardised gene-phenotype associations encoded by the Human Phenotype Ontology^8^ is another successful strategy that has led to the identification of disease genes by the phenotypic similarity of patients from different patient cohorts^9, 10^. Phenotype similarity analysis between model organisms and human disease patients has also highlighted new candidate gene-disease associations^11^. These successes led us to consider other features of genes that could be used to identify human disease genes.

Gene essentiality, or the requirement of a gene for an organism’s survival, is known to correlate with intolerance to variation^12^ and has been directly assessed in a number of species using high-throughput cellular and animal models^13–15^. In humans, essentiality has been investigated at the cellular level using cancer cell line screens based on gene-trap, RNAi or CRISPR-Cas9 approaches with the findings that ∼10% of protein-coding genes are essential for cell proliferation and survival^16–19^. The Broad Institute Project Achilles is extending this approach to characterise 1,000 cancer cell lines and, critically, corrects for copy number in their essentiality scoring^20, 21^. In parallel, the International Mouse Phenotyping Consortium (IMPC), a global research infrastructure that generates and phenotypes knockout (loss-of-function, LoF) mouse lines for protein coding genes, determines viability of homozygotes to assess gene essentiality^11, 22–24^. The number of observed homozygous LoF mice generated from an intercross between heterozygous parents allows the categorisation of a gene as lethal (0% homozygotes), subviable (<12.5% homozygotes) or viable, with ∼25%, ∼10% and ∼65% of genes classified as such, respectively^23^. These findings are consistent with results curated from the scientific literature indicating that approximately one-third of protein-coding genes are essential for organism survival^25^.

Several levels or definitions of essentiality are to be expected from these different model approaches^14^. “Core essential genes” are identified across different model systems, while other essential genes are dependent on context such as culture conditions, tissue or organism developmental stage^26–28^. Quantitative definitions of “low” and “high” gene essentiality have been proposed to account for the degree of dependency on external factors as well as the likelihood of a compensatory mutation rescuing necessary function^28^. Essential genes have also been classified by whether they are known to be associated with human disease, with functional mutations in non-disease-associated genes and with a mouse orthologue that is LoF embryonic lethal suggested as likely to prevent pregnancy, lead to miscarriage or to early death^29^. Other research has reported that orthologues of embryonic lethal LoF mouse genes are shown to have an increased association to diseases with high mortality and neurodevelopmental disorders^23, 30, 31^.

Here we provide a new <u>Fu</u>ll <u>S</u>pectrum of Intolerance to <u>L</u>oss-of-function (FUSIL) categorisation that functionally bins human genes by taking advantage of the comprehensive organismal viability screen performed by the IMPC and the cellular viability studies conducted by the Broad Institute Project Achilles. By exploring the FUSIL categories that span genes from those necessary for cellular survival all the way to those genes where loss of function has no detected phenotypic impact on complex organisms, we demonstrate a strong correlation of genes necessary for mammalian development with genes associated with human disease, especially early onset, multi-system, autosomal dominant disorders. Finally, we describe novel candidate genes for involvement in autosomal dominant, developmental disorders where potentially pathogenic variants had been identified in unsolved cases from three large-scale exome and genome datasets: Deciphering Development Disorders (DDD), 100,000 Genomes Project (100KGP) and the Centers for Mendelian Genomics (CMG) projects.

## RESULTS

### Cross-species comparisons divide genes into functional bins of essentiality/ viability

Previously, human cell-essential genes have been analysed^16–18^ in conjunction with lethal genes identified in the mouse^23^ by either characterising a core set of essential genes at the intersection^14^ or studying the union of the two^32^. In these studies, cellular essential genes were shown to have a nearly complete concordance with mouse lethal genes^19, 23^. Here, we have taken the human orthologue genes for which the IMPC has viability assessments and integrate the gene essentiality characterisation based on the cell proliferation scores determined by the Project Achilles Avana CRISPR-Cas9 screening performed on over 400 cell lines. This identified two sets of lethal genes: a set of cellular lethal genes essential for both a cell and an organism to survive (cellular lethal, CL), and a set of developmental lethal genes (DL) that are not essential at the cellular level but where LoF is lethal at the organism level. The IMPC viability pipeline also defines distinct sets of subviable (SV) and viable genes in LoF mice. The latter are further split into those with an abnormal phenotype (VP, viable with significant phenotype/s), or those with a normal phenotype (VN, viable with no significant phenotypes detected). As a result, we obtained 5 mutually exclusive phenotype categories reflecting the <u>Fu</u>ll <u>S</u>pectrum of Intolerance to <u>L</u>oss of function (FUSIL; Table 1). The correspondence between viable and SV genes in the mouse and non-essential genes in human cell lines was very strong (Table 1, Fig. 1a). An almost complete correspondence was also found between the mouse genes that are lethal in LoF strains and their human orthologues being essential in human cell lines. However, while 35% of genes lethal in the mouse were essential in human cell lines (CL bin), the remaining 65% have not been identified as cell essential and were classified as essential for organism development (DL bin). A near identical pattern was observed when other cellular essentiality datasets were used from previously published studies^16–18^, with most genes ending up in the same category (96% overlap; Supplementary Fig. 1c, Supplementary Table 1).

**Table 1.**
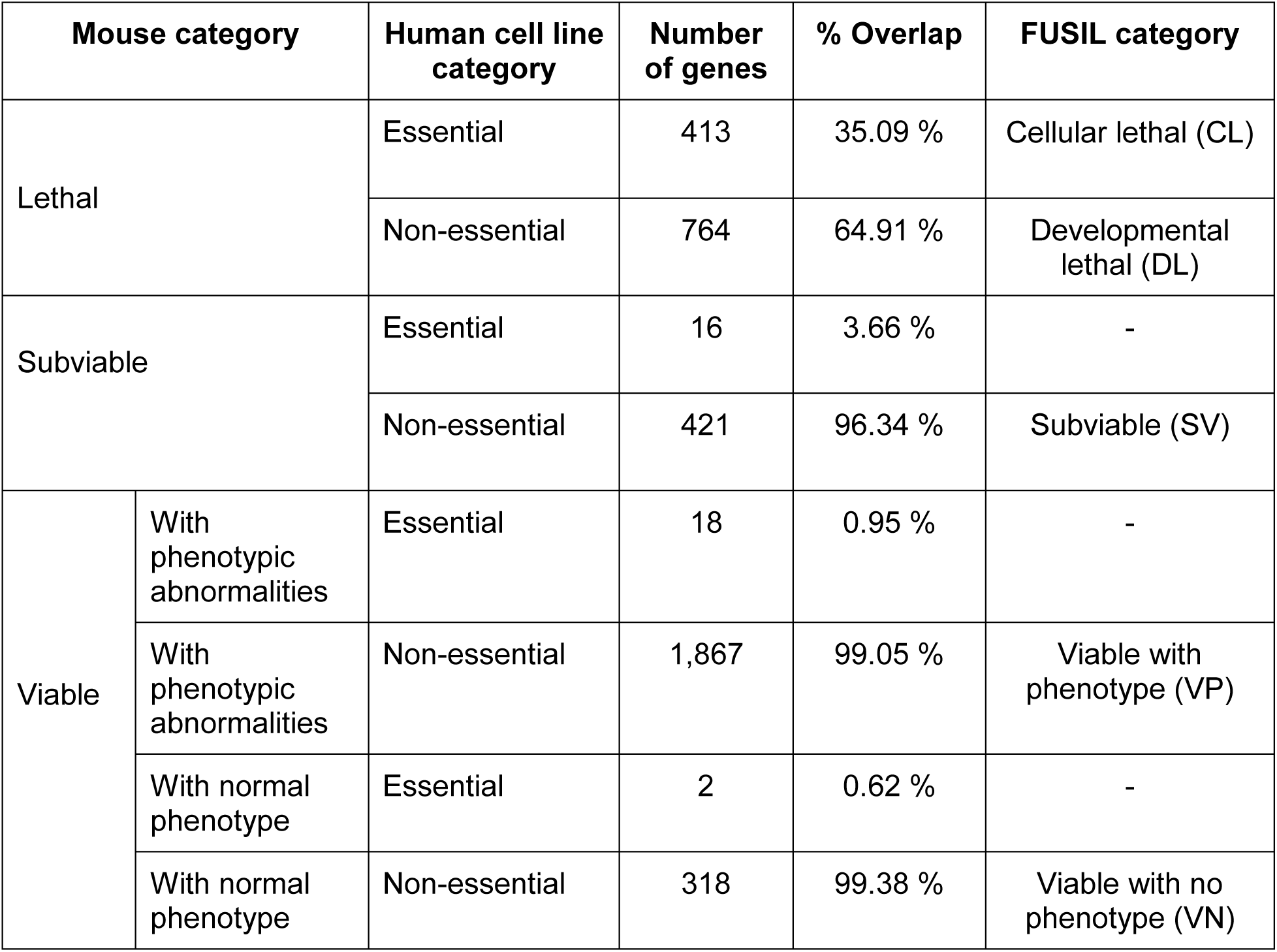
FUSIL categories. Integration of data from human cell essentiality screens from the Avana dataset and mouse phenotypes from IMPC screens defines 5 mutually exclusive categories of intolerance to loss of function.

| Mouse category |  | Human cell line category | Number of genes | % Overlap | FUSIL category |
| --- | --- | --- | --- | --- | --- |
| Lethal |  | Essential | 413 | 35.09 % | Cellular lethal (CL) |
|  |  | Non-essential | 764 | 64.91 % | Developmental lethal (DL) |
| Subviable |  | Essential | 16 | 3.66 % | - |
|  |  | Non-essential | 421 | 96.34 % | Subviable (SV) |
| Viable | With phenotypic abnormalities | Essential | 18 | 0.95 % | - |
|  | With phenotypic abnormalities | Non-essential | 1,867 | 99.05 % | Viable with phenotype (VP) |
|  | With normal phenotype | Essential | 2 | 0.62 % | - |
|  | With normal phenotype | Non-essential | 318 | 99.38 % | Viable with no phenotype (VN) |

**Fig. 1.**
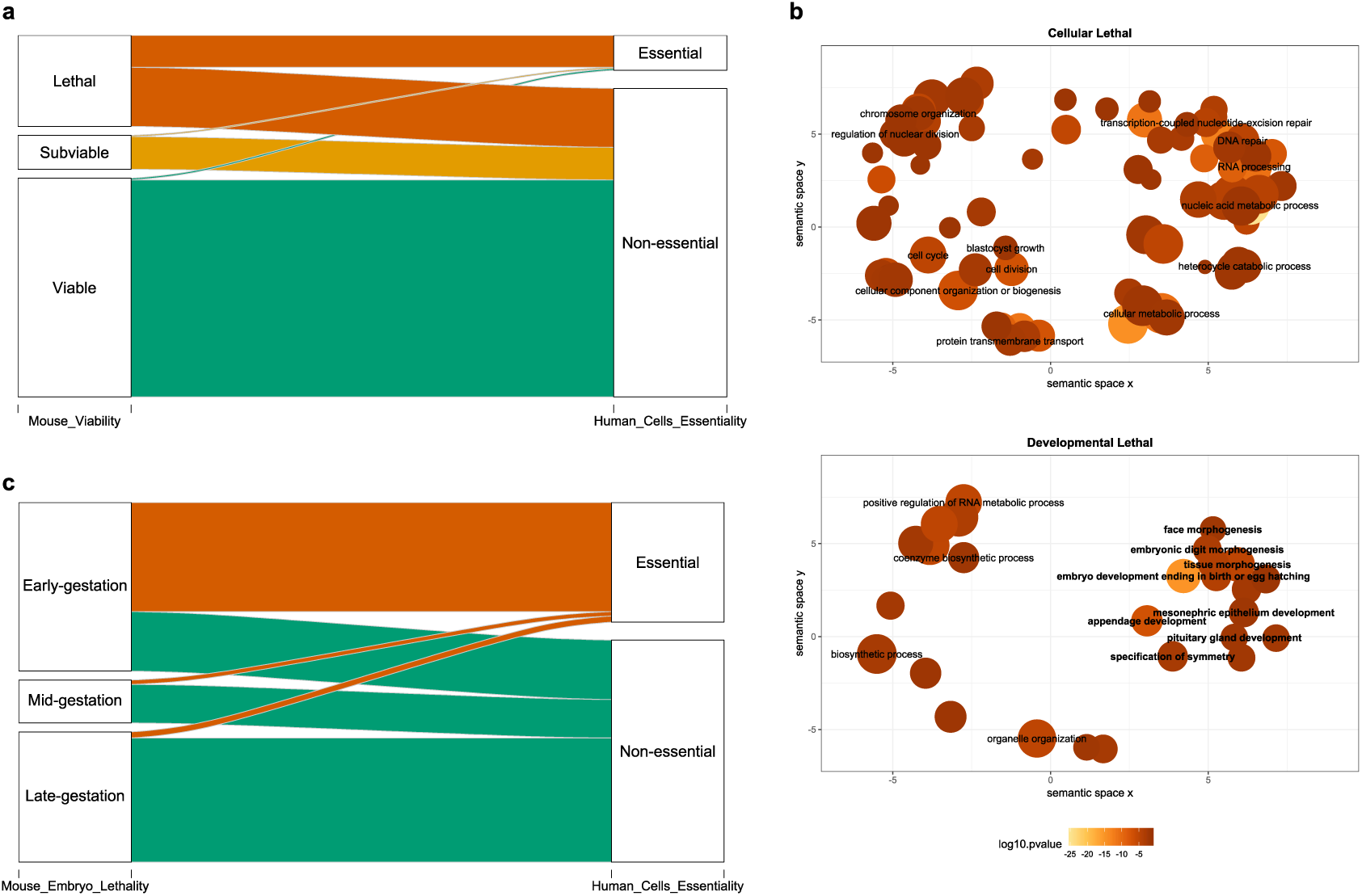
Cross-species FUSIL categories of intolerance to LoF. **a) Correspondence between primary viability outcomes in mice and human cell line screens.** The sankey diagram shows how human orthologues of mouse genes with IMPC primary viability assessment (lethal, subviable and viable) regroup into essential and non-essential human cell categories; the width of the bands is proportional to the number of genes. **b) Gene Ontology Biological Process (GO BP) enrichment results.** Significantly enriched GO terms at the biological process level were computed using the entire set of IMPC mouse-to-human orthologues as a reference (Table 1). Significant results were only found for the cellular and developmental lethal gene categories. Bubble size is proportional to the frequency of the term in the database and the colour indicates significance level as obtained in the enrichment analysis. The GO terms associated to embryo development are in bold. **c) Correspondence between mouse embryonic lethality stage and essentiality in human cell lines.** Embryonic lethal LoF strains are assessed for viability at selected stages during embryonic development: early (gestation) lethal (prior to E 9.5), mid (gestation) lethal (E9.5-E14.5/15.5), late (gestation) lethal (E14.5/E15.5 onwards). E, embryonic day.

### Genes essential for organism viability are involved in tissue development and morphogenesis

An enrichment analysis of Gene Ontology biological processes showed that the two lethal FUSIL categories (CL and DL) were clearly involved in different biological processes (Fig. 1b, Supplementary Table 2). Whereas the set of CL genes was enriched for nuclear processes (DNA repair, RNA processing, regulation of nuclear division, among other cellular processes), the DL genes were enriched in morphogenesis and development functions (such as embryo development, appendage development, tissue morphogenesis and specification of symmetry). In contrast, genes in the SV and viable categories (VP, VN) were not significantly enriched in any biological process despite reasonable sample sizes, probably reflecting diverse roles for these genes. These results provide additional evidence to make a distinction between the two sets of lethal genes in terms of their biological function.

### Early lethal genes in the mouse correlate with cellular essential genes

For those genes found to be lethal, the IMPC performed a secondary viability assessment of null homozygous embryos to determine the time of embryonic death. Three windows of lethality were classified for 400 lethal genes: early gestation (49.25%), mid-gestation (12.50%) and late gestation lethal genes (38.25%), confirming previous findings that nearly half of embryonic lethal mouse embryos die prior to embryonic day E9.5^23^. When this information was combined with the human cell dataset, we observed a strong concordance between the stage of embryo lethality and essentiality at the cellular level: 65% of early gestation lethal genes are essential in human cell lines, whereas only 10% of mid-gestation lethal genes and less than 5% of late gestation lethal genes fall into this category (Fig. 1c, Supplementary Table 3).

### Functional binning by viability and associated gene features

Essential genes have been shown to be located in regions with lower recombination rates^33, 34^ and we observe an increasing trend of recombination rates of the regions containing the genes across the FUSIL gene categories from most to least essential, with CL genes representing a clearly distinct category and still significant differences between bins for most pairwise comparisons (Fig. 2a, Supplementary Table 4). Higher expression values have also been previously associated with essential genes^16^ and here a decreasing expression, as measured by median GTEx expression across the entire range of tissues and cell lines, is observed from most to least essential FUSIL bins (Fig. 2b). Similar continuous trends were observed for other gene features previously associated with essential genes, including protein-protein interaction network properties (Fig. 2c) or the likelihood of the gene product being part of a protein complex (Fig. 2d). CL genes also stand out as a singular category regarding the number of paralogues (Fig. 2e).

**Fig. 2.**
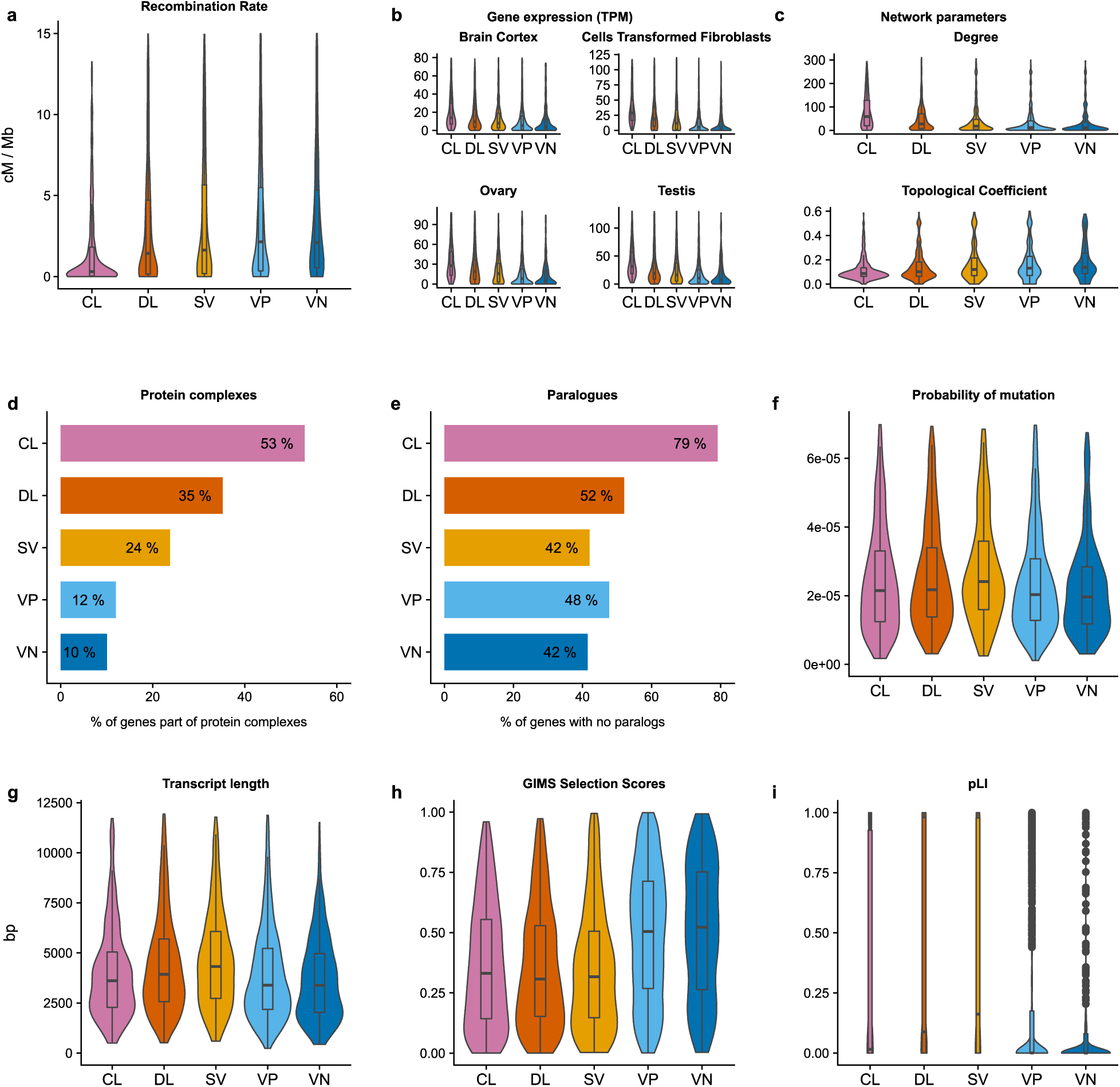
FUSIL categories and gene features. (CL, cellular lethal CL; DL, developmental lethal; SV, subviable; VP viable with phenotypic abnormalities; VN, viable with normal phenotype). **a) Violin plots showing the distribution of recombination rates for the different FUSIL bins.** Human recombination rates for genomic intervals^59^ were mapped to the closest gene and average recombination rates per gene were computed (outliers not shown). **b) Distribution of median gene expression values for different tissues.** Median TPM expression values from GTEx for selected non-correlated tissues are shown (outliers not shown). **c) Protein-protein interaction network parameters.** Violin plots showing the distribution of degree and topological coefficient computed from human protein-protein interaction data extracted from STRING. Only high confidence interactions, defined as those with a combined score of >0.7, were kept (outliers not shown). **d) Protein complexes.** Bar plots representing the percentage of genes in each FUSIL bin being part of a protein complex (human protein complexes). **e) Paralogues.** The barplot shows the percentage of genes without a protein-coding paralogue gene in each FUSIL bin. Paralogues of human genes were obtained from Ensembl Genes 95. A cutoff of 30% amino acid similarity was used. **f) Probability of mutation.** Distribution of gene-specific probabilities of mutation from Samocha, et al. ^67^(outliers not shown). **g) Transcript length.** Maximum transcript lengths among all the associated gene transcripts (Ensembl Genes 95, hsapiens dataset) (outliers not shown). **h) GIMS Selection Score.** Distribution of Gene-level Integrated Metric of negative Selection (GIMS)^68^ scores across the different FUSIL bins. **i) Probability of being Loss-of-Function intolerant scores (pLI)** retrieved from gnomAD2.1. Higher values indicate more intolerance to variation. Significance for pairwise comparisons for all features are shown in Supplementary Table 4.

In contrast, features associated with mutational rate appear to peak in the DL or SV bins: probability of mutation based on gene context (Fig. 2f), transcript length (Fig. 2g), and strength of negative selection measured by gene-level integrated metric of negative selection (GIMS) scores (Fig. 2h). However, these observed increases in the DL and SV genes relative to CL were only statistically significant for transcript length (Supplementary Table 4). A similar effect is observed with several intolerance to variation scores from large genomic studies, where higher (but not statistically significant) values are observed in the DL and SV genes relative to CL (Fig. 2i, Supplementary Fig. 2, Supplementary Table 4, Supplementary Table 5).

### Developmental lethal genes are enriched for human disease genes

Previous studies have reported associations between disease and essential genes, or just organismal essential genes, using different criteria and datasets^30, 35^. Our first report on developmental phenotypes showed a significant enrichment for disease genes in the IMPC essential genes^23^. By segmenting this set of genes into 3 mutually exclusive categories (CL, DL and SV; Fig. 3a), we found that whilst the CL and SV fractions showed a moderate enrichment for disease genes compared to all other categories (odds ratios, ORs > 1) and the two bins containing viable genes (VP and VN) were significantly depleted (ORs < 1), the highest overrepresentation of Mendelian disease genes was found in the DL fraction (2.6 fold-increased odds). This finding was consistent with an early study defining a set of peripheral essential genes^27^.

**Fig. 3.**
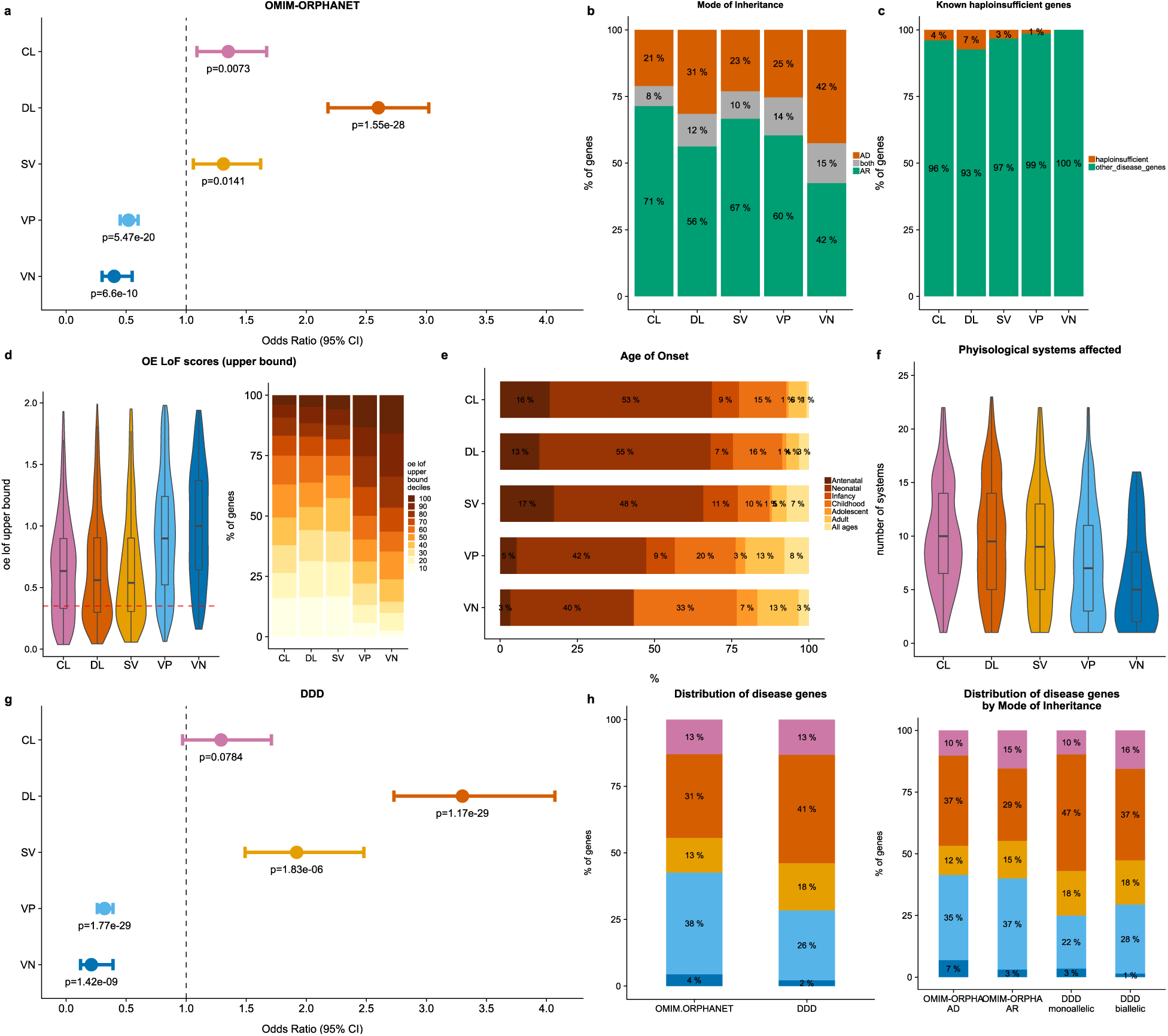
Human disease genes and FUSIL bins. (CL, cellular lethal CL; DL, developmental lethal; SV, subviable; VP viable with phenotypic abnormalities; VN, viable with normal phenotype). **a) Enrichment analysis of Mendelian disease genes**. Combined OMIM-ORPHANET data was used to compute the number of disease genes in each FUSIL bin. Odds Ratios were calculated by unconditional maximum likelihood estimation (Wald) and confidence intervals (CI) using the normal approximation, with the corresponding adjusted P-values for the test of independence. **b) Distribution of disease associated genes according to mode of inheritance**. Disease genes with annotations regarding the mode of inheritance according to the Human Phenotype Ontology^8^. **c) Haploinsufficient genes**. Known haploinsufficient genes curated by ClinGen (% with respect to the total number of disease genes in each bin). **d) Distribution of gnomAD o/e LoF scores**. Upper bound of observed versus expected LoF score (gnomAD 2.1.). A score threshold of 0.35 (dashed line) has been suggested to identify intolerant to LoF variation genes^57^. Distribution of genes across o/e LoF upper bound deciles. **e) Age of Onset** as described in rare diseases epidemiological data from Orphanet (Orphadata). The earliest age of onset associated to each gene was used. **f) Physiological systems affected**. The phenotypes (Human Phenotype Ontology, HPO) associated to each gene were mapped to the top level of the ontology to compute the number of unique physiological systems affected. **g) Enrichment analysis of developmental disorder genes**. DECIPHER^2^ set of genes was used to compute the number of developmental disorder genes in each FUSIL bin. **h) Distribution of disease genes**. Percent distribution of Mendelian and developmental disorder genes among the different FUSIL categories (colors representing FUSIL bins as in other figures). Percent distribution of disease genes by mode of inheritance.

Analysis of the mode of inheritance of the disease genes in each bin showed the CL fraction had the lowest proportion of autosomal dominant (AD) disorders, whilst the DL and VN fractions showed higher proportions (uncorrected P-values of 0.0143 and 0.004 respectively) (Fig. 3b), although the latter was based on a relatively small number of disease genes (40) with AD/AR annotations in this category relative to 119 and 286 seen in the CL and DL fractions respectively. The proportion of known haploinsufficient genes in the disease genes was greatest in the DL fraction with twice as many as in the CL fraction (Fig. 3c).

The Genome Aggregation Database (gnomAD) resource aggregates and harmonises exome and genome sequencing data from unrelated individuals representing broad populations and disease cohorts that are not known to be affected by a severe pediatric disease (http://gnomad.broadinstitute.org/). The analysis of the FUSIL categories for gnomAD’s observed/expected (o/e) LoF scores (upper bound fraction) which are used to evaluate a genés tolerance to LoF variation, showed that essential genes are more intolerant to inactivation compared to viable genes, and that the DL and SV categories showed the peak intolerance although the difference with respect to the set of CL genes was not significant (Fig. 3d, Supplementary Table 5). Again, this is consistent with an overrepresentation of AD disorders among DL genes. Similar results were found when we investigated other intolerance scores (Supplementary Fig. 2, Supplementary Table 5).

The proportion of disease genes associated with an early age of onset (antenatal/prenatal and neonatal) was highest in the CL and DL gene sets, with the percentage of later onset associated genes increasing as we move towards more viable categories (Fig. 3e). The degree of pleiotropy, measured by the number of physiological systems affected according to HPO annotations, followed a similar pattern, with CL and DL genes showing the highest number of affected systems (Fig. 3f).

In summary, our results suggest that disease genes in the DL fraction correlated with earlier ages of onset, multiple affected systems and autosomal dominant disorders (Supplementary Table 6). Given this, we compared the genes in our FUSIL bins with the genes associated with developmental disorders as reported by the DDD consortium in the Development Disorder Genotype - Phenotype Database (DDG2P) and found a strong enrichment in the DL FUSIL bin for developmental disease genes (3.3 fold-increase; Fig. 3g). The DDD resource has a larger representation of non-consanguineous patients with *de novo* mutations compared to the OMIM and Orphanet resources^36^, so we next compared enrichment of DL genes by mode of inheritance across the three resources and found the strongest enrichment for monoallelic disease genes in the DDD resource (Fig. 3h).

This result is particularly relevant for the identification of new disease genes using these FUSIL bins: for Mendelian genes, despite the enrichment in the DL fraction, the highest proportion was still found in the much larger VP bin (Fig. 3a, h), but for developmental disorders, as represented by DDG2P, the DL fraction contained the majority of genes, reaching a percentage close to 50% for monoallelic disease genes (Fig. 3g,h). Thus, we attempted to identify strong, novel candidate genes for undiagnosed cases of autosomal dominant, development disorders by extracting 163 genes from the DL bin genes (n=764) which had the following properties: (i) not described as associated with human disease by OMIM, Orphanet or DDG2P and (ii) highly intolerant to a LoF mutation (pLI > 0.90 or o/e LoF upper bound < 0.35 or HI < 10). We next focused on unsolved diagnostic cases from three large rare disease sequencing programs to investigate potential disease candidates within our set of 163 prioritised DL genes (Supplementary Table 7).

First, DDD makes publicly available a set of rare *de novo* and/or homozygous, hemizygous or compound heterozygous LoF research variants from unsolved cases of genetic developmental disorders in genes that are not associated with human disease according to OMIM and DDG2P, with nearly 14,000 children sequenced to date in the UK DDD study. Given recent findings from DDD that most developmental disorders cases could be explained by *de novo* coding mutations^36^ we searched for heterozygous, *de novo* variants in this dataset that affected any of the prioritised 163 DL candidate genes described above and found variants in 44 genes that met these criteria. Secondly, we searched 18,000 rare disease cases from the 100,000 Genomes Project (100KGP) and discovered *de novo* variants in undiagnosed patients with intellectual disability in 47 of the 163 genes in our candidate set.

Lastly, the Centers for Mendelian Genomics (CMG), a collaborative network of Centers to discover new genes responsible for Mendelian phenotypes provides a list of phenotypes studied and potential associated genes^37^ for some 2,000 genes. A set of 14 genes overlapping with the DL candidates are classified as either Tier 1 (mutations have been identified in multiple kindreds or fall within a significant linkage analysis peak or the phenotype has been recapitulated in a model organism) or Tier 2 genes (strong candidates but in most cases mutations within those genes have only been found in one kindred).

There was some degree of overlap between the candidates identified in the 3 programs (Fig. 4a) and for the next stage we focused on the genes with evidence from both the 100KGP (where we had detailed patient phenotypes and variant information) and either DDD (variants and high level phenotypes available) or the CMG (gene and high level phenotypes available). We particularly focused on 9 genes where the associated variants were not present in any population in gnomAD and each gene was also intolerant to missense variation (o/e missense < 0.8; Fig. 4b, Supplementary Table7). For these genes, further evidence for candidacy was gathered based on the number of unrelated families and phenotypic similarities between them, protein-protein interactions with known developmental disorder genes, embryonic and adult mouse gene expression in relevant tissues, and embryonic and adult mouse phenotypes that recapitulate the clinical phenotypes. Here we present two examples, *VPS4A* and *TMEM63B,* where the patient phenotype and genetic evidence is compelling as well as showing phenocopying in the mouse where the IMPC has produced the first knockout lines for these genes. For the other 7 genes, there was typically functional data strengthening the association but the patient evidence is currently less strong as we only have single, intellectual disability cases with detailed clinical phenotypes, or *de novo* variants are also observed in cases affected with other types of rare disease.

*VPS4A* (HGNC:13488, vacuolar protein sorting 4 homologue A) had no previously reported pathogenic variants and is highly intolerant to LoF and missense variants (gnomAD v.2.1., pLI = 0.928, o/e LoF = 0.139, o/e missense = 0.532)*. De novo* variants in *VPS4A* were detected in two unsolved, 100KGP intellectual disability cases but not in any of the other 18,000 cases representing most types of rare disease. These variants are not observed in gnomAD and both patients exhibited consistent intellectual disability, developmental delay, delayed motor development, microcephaly and eye abnormalities including cataracts (Supplementary Table 8, Supplementary Fig. 3). In addition, a Tier 2 CMG candidate was described with similar phenotypes of microcephaly, epilepsy, frontoencephalocele, right spastic hemiparesis and psychosocial retardation (Supplementary Table 8, Supplementary Fig. 3). The IMPC’s data for the first mouse knockout of the orthologous *Vps4a* gene, indicated preweaning lethality of the homozygotes and, in the case of the heterozygotes, abnormal skin morphology, enlarged spleen, and lens opacity, potentially modelling the eye phenotypes seen in the patients. LacZ staining in E12.5 embryos showed widespread expression. While the LoF mutants are lethal at P14, secondary viability of *Vps4a* mutants shows that they are viable at E18.5 (6/30, 20%), but display gross abnormalities at manual observation by and micro-Computed Tomography (CT) imaging. Homozygous *Vps4a* E18.5 embryos are smaller than wildtypes, with abnormal body curvature, omphalocele, small and compressed heart, abnormal spinal cord curvature, and abnormal brain development. Within the brain, microCT images showed evidence of abnormalities in the thalamus, thinning of the midbrain, and a smaller cerebellum and pons compared to the wild type (WT) littermates. The volume changes of the midbrain/ cerebellum/ pons might also be related to the enlargement of the fourth ventricle (Fig. 4c). Interestingly, *VPS4A* is known to directly interact in human with an intellectual disability gene, *CHMP1A,* from NMR, affinity chromatography, pull down and two hybrid assays, and both are part of the necroptosis and endocytosis pathways^38^. Variants in *CHMP1A* cause pontocerebellar hypoplasia type 8 (OMIM:614961), with similar phenotypes to patients with *VPS4A* variants: severe psychomotor retardation, pontocerebellar hypoplasia, decreased cerebral white matter, thin corpus callosum, abnormal movements, hypotonia, spasticity, and variable visual defects. GTEx shows particularly high levels of gene expression across all tissues, both disease and non-disease related. Similarly, high levels of expression were also seen in WT mouse embryos from 4 to 36 somites according to Deciphering the Mechanism of Developmental Disorders (DMDD)^39^.

**Fig. 4.**
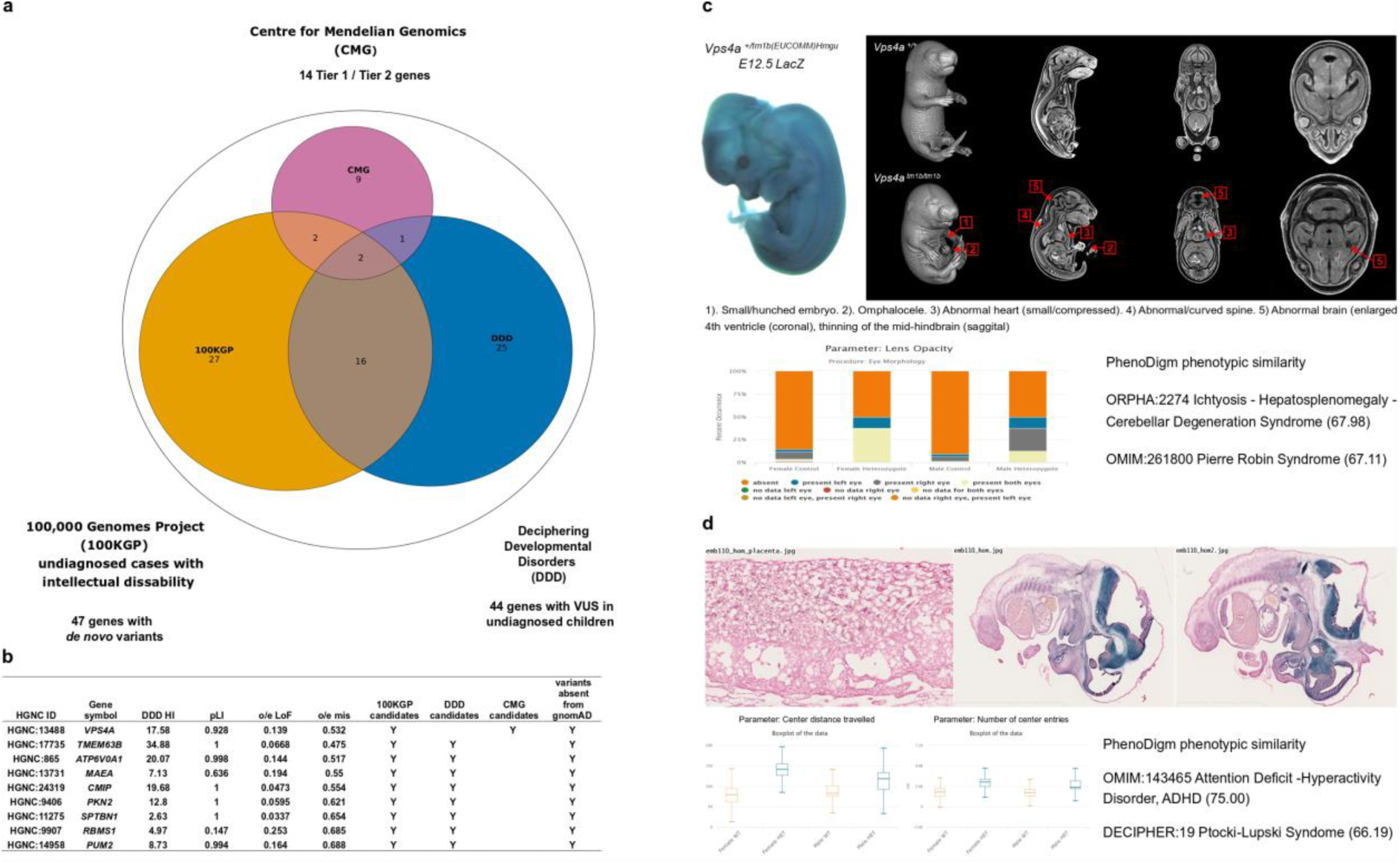
Developmental disorders gene candidate prioritisation. a) Venn diagram showing the overlap between DL prioritised genes with evidence from 3 large scale sequencing programs. Overlap between the set of 163 developmental genes highly intolerant to LoF variation (pLI > 0.90 or o/e LoF upper bound < 0.35 or HI < 10) and not yet associated to disease and the set candidate genes from three large rare disease sequencing consortia: 100KGP, CMG and DDD. **b) Set of 9 candidate genes**. The selected genes met the following criteria: (1) evidence from both the 100KGP (with detailed clinical phenotypes and variants) and either DDD (variants and high level phenotypes available) or CMG (gene and high level phenotypes available), (2) the associated variants were not present in gnomAD; and (3) intolerance to missense variation; these genes were further prioritised based on the number of unrelated probands and the phenotypic similarity between them and the existence of a mouse knockout line with embryo and adult phenotypes that mimic the clinical phenotypes. c) Mouse evidence for *VPS4A.* IMPC embryonic phenotyping of homozygous mutants at E18.5 showed abnormal/curved spine and abnormal brain among other relevant phenotypes. The phenotypic abnormalities observed in heterozygous knockout mice include lens opacity. Heterozygous mouse phenotypic similarity to known disorders as computed by the PhenoDigm algorithm. d) Mouse evidence for *TMEM63B.* IMPC homozygous mouse embryo lacZ imaging at E14.5 supporting neuronal expression during development. Heterozygous IMPC knockout mice associated phenotypes included abnormal behavior evaluated through different parameters. The heterozygous mice showed a high phenotypic similarity with several developmental disorder phenotypes.

*TMEM63B* (HGNC:17735, transmembrane protein 63B), is also extremely intolerant to loss of function and missense variants (gnomaD v. 2.1., pLI = 1.00, o/e LoF = 0.07, o/e missense = 0.475) but has no previously reported pathogenic variants. *De novo* variants in unsolved, developmental disorder cases were identified in one DDD case and four, unrelated 100KGP participants with intellectual disability but none of the other 18,000 100KGP cases (Supplementary Table 9). These variants are not observed in gnomAD and the exact same variant (ENST00000259746:c.130G>A) was detected in three of the families, causing a p.44Val>Met change in a transmembrane helix that is predicted to be pathogenic. The clinical data (Supplementary Table 9, Supplementary Fig. 4) available from the four 100KGP cases showed consistent intellectual disability and abnormal movement and brain morphology phenotypes, with seizures also observed in three of the patients. For the DDD case, the high level phenotypes available were consistent with the 100KGP cases. IMPC data for the first mouse knockout of the orthologous *Tmem63b* gene showed preweaning lethality of the homozygotes and, in the case of the heterozygotes, abnormal behaviour, hyperactivity and limb grasping phenotypes that are consistent with the human patients. Expression analysis placed *TMEM63B* in a cluster of genes that are expressed at medium levels from early to late development. GTEx showed high levels of gene expression in disease-related tissues, particularly for the brain cerebellum and muscular-skeletal tissues. High levels of expression were also seen across all mouse embryonic developmental stages (according to DMDD) with GXD data and the IMPC’s mouse embryo *lacZ* annotation supporting neuronal expression during development (Fig. 4d).

## DISCUSSION

The current diagnostic rate of large-scale, rare disease sequencing programs ranges from 20-40%^40, 41^ leaving the majority of patients without a diagnosis and with the associated personal, psycho-social and healthcare cost this entails. New disease-gene discovery methods are needed to complement deeper sequencing approaches currently being employed to identify disease-causal variants beyond single-nucleotide variants in the coding region^42^. Here we demonstrate that the FUSIL categorisation of gene essentiality, combined with intolerance to variation scores, patient phenotypes and their overlap with those observed in mouse lines with null alleles can assist in the prioritisation of disease-causal variant candidates. We show an enrichment for disease genes among developmental essential genes, which is consistent with the proposed model where disease associated genes occupy an intermediate position between highly essential and non-essential genes^29, 43, 44^. In this model, highly essential genes will not be associated with human diseases because any function-altering mutation will likely lead to miscarriage or early embryonic death. Our results provide evidence that highly essential genes needed for cellular processes are less likely to be associated with disease than developmental essential genes, suggesting a new complementary approach and resource for finding such disease genes and understanding disease mechanisms.

An interesting finding was the dichotomy of trends observed for gene-associated features. For certain gene features, we replicate previously observed trends where genetic features are most differentiated between the two ends of the FUSIL spectrum e.g. genes with paralogues, gene expression, number of protein-protein interactions or the likelihood of being part of a protein complex^16, 29, 45^. The CL bin shows the lowest rates of recombination, and both the CL and DL fractions exhibit significantly lower rates than the viable categories. This is consistent with previous findings that genes with essential cell functions, and showing accumulation of disease-associated mutations, concentrate in genomic regions with suppressed recombination^33, 34^. The strong enrichment of CL genes for presence in protein complexes and a lack of paralogues would suggest these genes should be particularly intolerant to damaging mutations with no functional compensation that has evolved to buffer critical cell processes^16, 46, 47^. For other gene features associated with mutational rate, the trends peak in the DL and SV bins (e.g. transcript length, constraint scores) leading to the counter-intuitive observation that CL genes are less constrained against variation than DL genes, however the differences between these two categories are non-significant for the different constraint scores evaluated. The significant enrichment for longer transcript lengths and Mendelian disease genes aligns with previous observations that disease genes tend to be longer^48, 49^, and genetic variant intolerance metrics such as shet, pLI, and o/e LoF are highly dependent on gene length^7, 14^. In fact, one caveat of these scores is that short genes and recessively acting genes may go undetected^14^.

Recent analysis from the DDD study estimates that *de novo* heterozygous variants will be the cause for nearly half of all undiagnosed, non-consanguineous individuals with developmental disorders^36^. Since these early onset disorders are particularly overrepresented among developmental lethal genes, we focused on a set of unsolved developmental disorder cases from DDD, the 100KGP and CMG. Research candidates were identified in half (82) of the 163 highly LoF intolerant genes we selected from the DL fraction that have not previously been associated with Mendelian disease. Nine genes in particular were observed in multiple consortia and are based on *de novo* variants in unsolved cases of developmental disorders that have never been observed in the general population. Further exploration of overlapping genotypes and phenotypes in the patients as well as embryonic and adult mouse phenotype evidence from the IMPC, alongside expression and protein-protein interactions with known developmental disorder genes, strengthened these novel associations. Two genes were particularly compelling from a clinical perspective as variants were only observed in intellectual disability cases and in multiple, unrelated families with specific phenotypes in common. The same *de novo* variant in *TMEM63B* was observed in 3 separate DDD and 100KGP patients with consistent intellectual disability, movement and brain morphology phenotypes that were recapitulated in the IMPC *Tmem63b* mutant mouse. Concordant phenotypes were also observed in 3 patients with *VPS4A* variants from the CMG and 100KGP and in the E18.5 mutant mouse embryos from the IMPC. Future identification of other families with similar variants segregating with disease and functional characterisation of the specific human variants will be required to establish a definitive role for these genes in disease. The IMPC partners are already supporting the CMG, 100KGP, other disease sequencing projects such as KidsFirst, and the wider rare disease community through Crispr/Cas9 production and phenotyping of mouse lines modelling patient-specific, potentially pathogenic variants.

Cross-species data integration is not without its limitations. Not all human genes have a clear one-to-one orthologue in the mouse genome; in particular certain gene families, such as immune receptor genes, are rapidly evolving in both species. In addition, a significant proportion of the mouse protein-coding genome is not yet phenotyped by the IMPC and is without viability data. We chose to focus on high-quality, public IMPC viability calls based on robust statistical methodologies in this investigation, and not to integrate the numerous, additional data from literature curation of mice with knockout alleles because the variation in methods, genetic context and gene targeting approach are known to affect embryonic lethality. Indeed, we observed a ∼10% disagreement between embryonic lethality published in the literature on a variety of genetic backgrounds *vs* the IMPC observations made exclusively on an inbred C57BL/6N background (Methods and Supplementary Fig. 5). Human cell line data also has caveats, including the haploid nature of some of the cell lines and the use of immortalised cell lines, which may result in the identification of some genes as essential that are not in the context of multi-cellular systems. The gene-based constraint scores based on human sequencing data primarily identify selection against heterozygous variation and may fail to detect short genes and recessively acting genes, while the homozygous viability screens in knockout mice typically measure recessive effects. In fact, for 1,015 genes that were lethal in homozygous LoF mouse lines and further investigated in heterozygous mice, 1,013 resulted in viable phenotypes. Hence, a moderate overlap between the set of essential genes identified through the different approaches is to be expected^14^. Further investigation into haploinsufficient, essential genes in the mouse is more technically challenging but may reveal further disease candidates.

In summary, this study highlights how the information on gene essentiality may be used to prioritise potential pathogenic variants in new disease genes from human sequencing studies. Clinical researchers assessing candidate disease genes should consider using high quality model organism data in conjunction with gene constraint scores from human sequencing projects. We intend to incorporate such data into our Exomiser variant prioritisation tool^50^, which together with patient phenotypes, may facilitate the genetic diagnosis by prioritising genes in the relevant FUSIL category. Large-scale projects such as the IMPC and Broad Institute Project Achilles are continuing to generate ever larger datasets, making the resources richer and more robust for these analyses. Future work will explore the mechanisms behind how these redefined essential categories correlate to other functional attributes and, ultimately, the evolutionary constraints imposed upon gene essentiality.

## METHODS

In this study, we examined genes for which viability data for null homozygote mice has been produced by the International Mouse Phenotypic Consortium (IMPC, www.mousephenotype.org). We obtained high- to moderate-confidence human orthologues and integrated selected human genetic data. In particular, we incorporated gene viability data obtained in human cell screens (initially 11 cell lines corresponding to three studies)^19^; in this study we used the Broad Institute Project Achilles Avana dataset comprising 485 cell lines, release 18Q3 of August 2018). We also incorporate constraint scores (shet, RVIS, pLI and O/E LoF), disease information (OMIM, ORPHANET, HPO, DDD) and DMDD annotations (see below for details). We use HGNC and MGI as stable identifiers in our analysis to avoid problems associated with gene symbol changes (synonyms and previous symbols, which may lead to ambiguous identifiers).

### IMPC primary and embryo viability assessment

We conducted an analysis on 4,934 genes with primary adult viability data currently available in IMPC Data Release (DR) 9.1. This release included the 1,751 genes previously analysed in Dickinson, et al. ^23^ and the 4,237 genes analysed in Munoz-Fuentes, et al. ^19^, corresponding to DR4.0 and DR7.0, respectively, and additional data collected since then. Viability data generated by the IMPC was analysed as defined in IMPRESS (the International Mouse Phenotyping Resource of Standardised Screens, https://www.mousephenotype.org/impress/). A minimum of 28 pups were genotyped before weaning, and the absence of knockout (null) homozygote pups would classify the gene as lethal. Thus, lethal lines are defined as those with an absence of live null homozygous pups, while subviable lines are those with fewer than 12.5% live homozygous pups (half of the 25% expected; *P* < 0.05, binomial distribution). Viable mouse lines are those for which homozygous (null and wild type) and heterozygous pups are observed in normal Mendelian ratios. A viable call was also made when there were less than 28 total pups and homozygous null pups ≥ 4 (as this would result in ≥ 14% homozygous pups when 28 pups were genotyped). We filtered out genes for which sample size was insufficient (total pups < 28, *n* = 1 gene), hemizygous genes (*n* = 13 genes), as well as those with conflicting calls (genes that appear in more than one viability category, *n* = 34 genes). The resulting set comprises 4,886 genes, of which 1,171 had a lethal phenotype, 449 a subviable phenotype, and 3,266 a viable phenotype (24%, 9% and 67%, respectively).

The IMPC also implements a dedicated embryonic pipeline for lethal lines, in which null homozygous embryo viability is assessed at selected stages during embryonic development, including embryonic day (E) 9.5, E12.5, E14.5-E15.5, and E18.5. Viability at a given stage is assessed by scoring homozygous embryos for the presence of a heartbeat at dissection, as described in Dickinson, et al. ^23^. To establish windows of lethality, we considered a gene lethal at a given stage if no live homozygous embryos were identified after scoring at least 28 live embryos, and viable if any live homozygous embryos were identified, irrespective of additional phenotype features. Following Dickinson, et al. ^23^, we defined windows of lethality as “prior to E9.5”, “E9.5-E12.5”, “E12.5-E14.5/E15.5”, “E15.5-E18.5”, “after E14.5/E15.5”, and “after E18.5”. Lines with incomplete data to define these windows were excluded. For this study, windows were combined to yield three functional groups: early (gestation) lethal (prior to E9.5), intermediate (mid gestation) lethal (E9.5-E12.5, E12.5-E14.5/E15.5) and late (gestation) lethal (E14.5/E15.5-E18.5, after E14.5/E15.5 and after E18.5). Out of 523 for which secondary screen data are available, 400 could be assigned to one of the described windows. Additional time points are required to complete the window assignment for the remaining 123 genes (Supplementary Table 3).

### Orthologue mapping

Orthologues were obtained using the HCOP tool developed by HGNC, based on 12 established inference methods (https://www.genenames.org/tools/hcop/, HCOP file with human and mouse orthologue inferences downloaded 18.10.31)^51^. We determined the confidence on the orthologue prediction based on the number of methods that supported each inference (12-9 methods, 100-75%, good-confidence orthologue; 8-5, 67-42%, moderate-confidence orthologue; 4-1, 33-8%, low-confidence orthologue; 0 - no orthologue). We kept orthologues for which at least one gene, the human or the mouse gene, was protein coding and the orthologue inference score was ≥ 5. Of those, we kept genes for which in both directions, mouse-to-human and human-to-mouse, the score was maximum (and also filtered out genes with duplicated maximum scores). This resulted in 4,664 genes. Of these, 33 genes with conflicting viability phenotypes and 17 genes with no adult viability data and insufficient embryo data to call a phenotype were not considered further. Among the remaining 4,614 genes, 1,185 (26%) genes had a lethal phenotype, 443 (9%) a subviable phenotype, and 2,986 (65%) a viable phenotype. Of these, 4,446 genes had human cell viability data (Avana data set; see next section).

### Human cell essentiality

In our previous study^19^, we used viability data as reported for 11 cell lines from 3 studies^16–18^. Here we used the Broad Institute Project Achilles Avana dataset, CRISPR-Cas9 proliferation (essentiality) scores. This data set comprises viability data on 17,634 genes in 485 cell lines (release 18Q3 in August 2018), with lower values indicating more intolerance (more essential). The Avana data set presents several advantages as compared to the previous studies. It is a larger data set, viability is measured using the same units for all cell lines (allowing us to obtain a mean per gene) and the data are corrected for copy number variation^21^. Gene identifiers were provided as Entrez identifiers (NCBI) and converted to HGNC identifiers.

### Determining functional bins

For each gene, we obtained a mean of the Avana proliferation scores and integrated with our dataset of mouse genes with viability data based on the mouse-to-human orthologues (obtained as described above). We observed that for genes with an Avana mean score below -0.45, the mouse null homozygotes were lethal in almost all cases, while genes with an Avana mean score above -0.45 presented lethal, subviable or viable phenotypes (Supplementary Fig. 1a, Supplementary Fig. 1b). A similar pattern was observed when a different resource for cell essentiality –based on 11 cell lines from 3 different studies – was used (methods and threshold criteria explained in Munoz-Fuentes, et al. ^19^ (Supplementary Fig. 1c).

F1 scores were derived from confusion matrices generated when considering different Avana mean scores and the classification from the previous studies, and a mean score cut-off of -0.45, was found to maximise the F1 scores across the different datasets (Supplementary Fig. 1d, Table 1, Supplementary Table 1). We therefore set a threshold where a mean Avana score equal to or below -0.45 were considered essential in cell lines, while the set above -0.45 corresponded to cellular non-essential genes. Based on all this evidence, we categorised two sets of genes, Cellular Lethal (CL) and Developmental Lethal (DL) genes, comprising 413 and 764 genes, respectively (Table 1).

Genes with a subviable or viable phenotype were classified for their human orthologue almost in all cases as non-essential (Avana mean score for each gene > -0.45), except for 16 and 22 genes for the subviable and viable categories, respectively, which had a mean Avana scores below but very close to the -0.45 threshold). These genes were non-essential based on data from the 3 human cell studies. Thus, we distinguish a Subviable group (SV) with Avana mean score > -0.45, as well as two outlier groups, SV.outlier and V.outlier, with Avana mean score ≤ - 0.45. We classified viable genes further, in those with at least one IMPC significant phenotype (VP) or no IMPC significant phenotypes (VN). A number of viable genes (n=627) had less than 50% phenotype procedures obtained so far as the IMPC releases are snapshots of ongoing characterisation for many lines, and thus we termed them V.insuffProcedures (Supplementary Table 10).

These categories, established using the Avana cell scores, were almost identical (96% concordance) to those established using a gene essentiality classification (essential / non-essential) based on the 11 cell lines from 3 studies as described previously^19^.

All the subsequent analysis focused on the main five FUSIL bins: CL, DL, SV, VP and VN. Sankey diagrams representing the mappings between mouse and human cell essentiality categories were plotted with the R package *alluvial*^52^. Gene Ontology Biological Process enrichment was conducted with the R package *category*^53^, with the entire set of IMPC genes as the reference set. BH method was applied for multiple testing correction^54^ and plotted using the REVIGO algorithm for semantic similarity and redundancy reduction^55^. The algorithm selects GO term representatives out of the significant results, maximising semantic representation and enriched/statistical significance (settings: SimRel semantic similarity measure, medium similarity, Homo sapiens database). Significant results were only found for CL and DL gene categories. In Fig. 1b, bubble size is proportional to the frequency of the term in the database and the colour indicates significance level as obtained in the enrichment analysis

### Previous knowledge on mouse viability

The IMPC is an ongoing project. Given that about ¼ of the mouse genome has been screened for viability to date, we decided to compare our results with previous annotations from Mouse Genome Informatics (MGI)^56^. We found a total of 4,599 mouse genes with embryo lethality annotations (50 Mammalian Phenotype Ontology terms as described in Dickinson, et al. ^23^ once conditional mutations were excluded from a total of 13,086 genes with any phenotypic annotation (including normal phenotype). That would mean that approximately 35% of the genes for which a knockout mouse has been reported in the literature and curated by the MGI showed some type of embryo / perinatal lethality. As the MGI resource also includes IMPC data, we subsequently excluded those gene-phenotype associations corresponding to IMPC alleles. This results in 9,254 mouse genes with phenotypic annotations other than IMPC with a PubMed ID (old papers with no abstract available, conference abstracts and direct data submissions were therefore excluded for this analysis). For 3,362 (36%) of these genes, we found annotations related to embryonic lethality.

Even though the procedures for determining viability may differ from the standardised viability protocol followed by the IMPC, this percentage is very close to that found using IMPC data, with 26% of the screened genes lethal and 10% of the screened genes subviable. For 2,115 mouse genes with both IMPC and non-IMPC phenotypic annotations available to infer viability, we found discrepancies for a set of 63 genes that were scored as lethal by the IMPC and with no previous records of lethality in MGI, as well as 154 genes scored as viable by the IMPC and with some type of lethality annotation reported in MGI (Supplementary Fig. 5, Supplementary Table 11). (Mouse Genome Database (MGD) at the Mouse Genome Informatics website [http://www.informatics.jax.org; MGI_GenePheno.rpt; Data Accessed 19.02.06].

### Constraint scores of gene variation based on human population data

The Residual Variation Intolerance Score (RVIS)^5^ was downloaded from http://genic-intolerance.org/, version CCDSr20, with lower values indicating more intolerance to variation. Estimates of selection against heterozygous protein-truncating variants (shet) were obtained from the supplementary material of Cassa, et al. ^12^ with higher values indicating more intolerant to variation. The probability of a gene being intolerant to loss of function (pLI)^6^, with higher values indicating more intolerance, the observed / expected (o/e) ratio of Loss of Function (LoF) with the corresponding upper bounds (LOEUF) the (o/e) ratio of missense and the (o/e) ratio of synonymous alleles, with lower values indicating more intolerance to variation, were retrieved from gnomAD2.1 (http://gnomad.broadinstitute.org)^7,57^. The haploinsufficiency score (HI), was obtained from the Deciphering Developmental Disorders (DDD) consortium (https://decipher.sanger.ac.uk/about#downloads/data, downloaded BED file, 18.11.27, Haploinsufficiency Predictions Version 3)^58^. High ranks (e.g. 0-10%) indicate a gene is more likely to exhibit haploinsufficiency while low ranks (e.g. 90-100%) indicate a gene is more likely to NOT exhibit haploinsufficiency. In all cases, gene identifiers were obtained as symbols and converted to HGNC IDs using the multi-symbol checker provided by HGNC (https://www.genenames.org/tools/multi-symbol-checker/).

### Gene features

#### Recombination Rates

We used the average genetic map computed from the paternal and maternal genetic maps from Halldorsson, et al. ^59^ (Data S3). The recombination rate (cMperMb) in intervals provided were mapped to the closest protein coding gene - Ensembl 96 GRCh 38 - (upstream or downstream) by means of R_bedtools_closest function in *HelloRanges* library^60^. Gene positions were obtained through *biomaRt*^61^. Once we assigned the recombination rates for the intervals provided to a certain gene, average recombination rates per gene were computed.

#### Gene expression

Gene median TPM values by tissue were downloaded from the GTEx portal [https://gtexportal.org/home/datasets, Accessed 18.12.02, file” GTEx_Analysis_2016-01-15_v7_RNASeQCv1.1.8_gene_median_tpm.gct.gz”]^62^. Gene symbols were mapped to HGNC ID identifiers. Spearman correlation coefficients between gene expression values across tissues were estimated, only TPM values for relevant (non-correlated) tissues were considered for further analysis.

#### Protein-protein interaction data

Human protein-protein interaction data were downloaded from STRING website [Accessed 18.11.18]. Only high confidence interactions, defined as those with a combined score > 0.7 were used for further analysis^63^. Ensembl protein IDs were mapped to HGNC IDs using Ensembl biomaRt (Ensembl Genes 94)^61^. Several network parameters for every node in the network were computed with Cytoscape (v3.5.1) plug-in Network Analyzer^64, 65^. Spearman correlation coefficients between different network parameters were estimated, only non-correlated parameters were considered for further analysis.

#### Protein complexes

A core set of human protein complexes was downloaded from Corum 3.0^66^ (http://mips.helmholtz-muenchen.de/corum/#download) [Accessed 18.12.03]. Gene symbols were mapped to unambiguous HGNC ID identifiers corresponding to protein coding genes.

#### Paralogues

Paralogues were retrieved with biomaRt (https://www.ensembl.org/biomart). Ensembl Genes 95 version [Data accessed 19.02.18]. Only genes with HGNC IDs were considered, and only those protein coding paralogues with an HGNC ID were kept for downstream analysis. A cut-off of 30 for the percent of identical amino acids in the paralogue compared with the gene of interest was used for the computation.

#### Probability of mutation

Per-gene probabilities of mutations (all types of mutations) were obtained from Samocha, et al. ^67^. Gene symbols were mapped to HGNC IDs. (Probabilities are shown as 10∧all).

#### Transcript length

Transcript lengths were retrieved with biomaRt R package (Ensembl Genes 96, hsapiens_gene_ensembl dataset). For each HGNC ID, the maximum transcript lengths was computed from all the gene transcripts^61^.

#### Selection Scores

Gene-level Integrated Metric of negative Selection (GIMS) scores, which combine multiple comparative genomics and population genetics to measure the strength of negative selection were obtained from Sampson, et al. ^68^ (Table_S1) and Alyousfi, et al. ^69^. Gene symbols were mapped to unambiguous HGNC ID identifiers.

### Human disease genes annotations and analysis

#### Gene-disease associations

Disease associated genes curated by OMIM^70^ and Orphanet^71^ were analysed through our PhenoDigm pipeline^72^ [Data accessed 19.01.16]. DECIPHER developmental disorders genes are defined as those “reported to be associated with developmental disorders, compiled by clinicians as part of the DDD study to facilitate clinical feedback of likely causal variants. The DDG2P is categorised into the level of certainty that the gene causes developmental disease (confirmed or probable), the consequence of a mutation (loss-of function, activating, etc) and the allelic status associated with disease (monoallelic, biallelic, etc)” [DDG2P version: DDG2P_19_2_2019.csv; https://decipher.sanger.ac.uk/ddd#ddgenes].

#### Mode of inheritance and physiological systems affected

Mode of Inheritance and number of physiological systems affected were annotated for each gene according to Human Phenotype Ontology annotations [https://hpo.jax.org/app/download/annotation, Downloaded 19.02.19]. The file ALL_SOURCES_ALL_FREQUENCIES_genes_to_phenotype.txt provides a link between genes and HPO terms^8^. Autosomal recessive inheritance (HP:0000007) and Autosomal dominant inheritance (HP:0000006) annotations were selected for downstream analysis. Phenotype ontology terms associated with each gene were mapped to the top level of the HPO to compute the number of unique physiological systems affected.

An additional set of haploinsufficient genes from ClinGen^73^ (https://www.clinicalgenome.org/) (n=295) was used for the analysis [https://github.com/macarthur-lab/gene_lists; Data accessed 19.02.19].

#### Age of onset

The age of onset was obtained from rare diseases epidemiological data (Orphadata) [http://www.orphadata.org/cgi-bin/epidemio.html; Data accessed,19.02.13]. The earliest age of onset associated with each gene was selected for downstream analysis.

#### Disease gene enrichment

For each FUSIL bin, odds ratios were computed from a contingency table with the number of disease (combined OMIM and ORPHANET) and non-disease genes for each one of the categories versus the remaining set of IMPC genes with FUSIL information. Odds Ratios were calculated by unconditional maximum likelihood estimation (Wald) and Confidence Intervals using the normal approximation, with the corresponding two-sided P-values for the test of independence calculated using Fisher’s exact test as implemented in the R package *epitools*^74^ (adjusted P-values, BH adjustment^54^).

#### Candidate developmental disorder genes annotation and prioritisation strategy

A gene set consisting of those developmental lethal genes (n = 764) that were not associated to a Mendelian disorder according to OMIM, ORPHANET or DECIPHER (n=387) and highly likely to be haploinsufficient (HI % < 10 | o/e lof upper bound < 0.35 | pLI > 0.90) (n=163) was used to identify candidate genes for undiagnosed cases of developmental disorders with heterozygous mutations.

DECIPHER developmental disorders (DDD)^2^ research variants (found in ∼2,000 genes) were downloaded from https://decipher.sanger.ac.uk/ddd#research-variants [Accessed 18.11.05]. Centers for Mendelian Genomics (CMG)^75^ Tier 1 and Tier 2 level genes (∼ 2,000 genes) were obtained from http://mendelian.org/phenotypes-genes [Accessed 19.28.02]. *De novo* variants in undiagnosed patients with intellectual disability from the 100,000 Genomes Project (100KGP) and associated clinical phenotypes (HPO terms) were extracted by querying the data available in the GeCIP research environment [Accessed 18.10.20] and intersecting with our set of 163 prioritised genes. Research candidates were identified in 82 of the 163 prioritised genes (highly LoF intolerant not previously been associated with Mendelian disease). Out of the total number of 47 genes with heterozygous *de novo* variants from undiagnosed cases in the 100KGP project and with extensive clinical phenotype available, 19 overlap with genes with heterozygous *de novo* variants from the DDD set of research candidates and 4 of them with Tier1 or Tier 2 genes from CMG (of which 2 were shared between the 3 datasets). We next focused on this set of overlapping genes to narrow down the search for strong candidates, we discarded those genes where the variants were present at any frequency in gnomAD along with those intolerant to missense variation (gnomAD o/e missense score < 0.8). This resulted in 9 genes that were then prioritised based on the presence of unrelated probands with phenotypic similarities and the existence of knockout mice – with embryonic and/or adult phenotypes - mimicking the clinical phenotypes.

The R package *eulerr*^76^ was used to create the Venn diagram and the *ontologyPlot*^77^ R package was used for the visualisation of the patient phenotypes as subgraphs of the HPO. For the set of candidate genes, expression analysis was conducted using BrainSpan^78^ and FUMA Gene2Func^79^ with all protein-coding genes as a background. Other mouse annotation resources include: Deciphering the Mechanism of Developmental Disorders (DMDD)^39^ (https://dmdd.org.uk/); GXD resource^80^; IMPC mouse embryo lacZ imaging (http://www.mousephenotype.org/data/imageComparator?&parameter_stable_id=IMPC_ELZ_063_001&acc=MGI:2387609).

#### Software

R software^81^ including the following additional packages were used for data analysis and visualisation: *dplyr*^82^, *ggplot2*^83^, *cowplot*^84^, *ggpubr*^85^.

#### Ethical approval

Mouse production, breeding and phenotyping at each centre was done in compliance with each centre’s ethical animal care and use guidelines in addition to the applicable licensing and accrediting bodies, in accordance with the national legislation under which each centre operates. All efforts were made to minimize suffering by considerate housing and husbandry. All phenotyping procedures were examined for potential refinements disseminated throughout the IMPC. Animal welfare was assessed routinely for all mice.

All patient data used in this study was either accessed through the public websites provided by DDD and the CMG or, in the case of the 100KGP, through the research environment provided by Genomics England and conforming to their procedures. All participants in the 100KGP have provided written consent to provide access to their anonymised clinical and genomic data for research purposes.

## Acknowledgements

This work was supported by NIH grant U54 HG006370. IMPC-related mouse production and phenotyping was funded by the Government of Canada through Genome Canada and Ontario Genomics (OGI-051) for NorCOMM2 (C.M.) and the National Institutes of Health and OD, NCRR, NIDDK and NHLBI for KOMP and KOMP2 Projects U42 OD011175 and UM1OD023221 (C.M., K.C.K.L), Infrafrontier grant 01KX1012, EU Horizon2020: IPAD-MD funding 653961 (M.H.d.A); EUCOMM: LSHM-CT-2005-018931, EUCOMMTOOLS: FP7-HEALTH-F4-2010-261492 (W.G.W). UM1 HG006348; U42 OD011174; U54 HG005348 (A.L.B), NIH U54706HG006364 (A.L.B). Wellcome Trust grants WT098051 and WT206194 (D.A). The French National Centre for Scientific Research (CNRS), the French National Institute of Health and Medical Research (INSERM), the University of Strasbourg and the “Centre Europeen de Recherche en Biomedecine”, and the French state funds through the “Agence Nationale de la Recherche” under the frame programme Investissements d’Avenir labelled (ANR-10-IDEX-0002-02, ANR-10-LABX-0030-INRT, ANR-10-INBS-07 PHENOMIN (J.H.). This research was made possible through access to the data and findings generated by the 100,000 Genomes Project. The 100,000 Genomes Project is managed by Genomics England Limited (a wholly owned company of the Department of Health). The 100,000 Genomes Project is funded by the National Institute for Health Research and NHS England. The Wellcome Trust, Cancer Research UK and the Medical Research Council have also funded research infrastructure. The 100,000 Genomes Project uses data provided by patients and collected by the National Health Service as part of their care and support and we are grateful to both for making this available. We are also grateful for the data access provided by the DDD and CMG projects. The DDD study presents independent research commissioned by the Health Innovation Challenge Fund [grant number HICF-1009-003], a parallel funding partnership between Wellcome and the Department of Health, and the Wellcome Sanger Institute [grant number WT098051]. The views expressed in this publication are those of the author(s) and not necessarily those of Wellcome or the Department of Health. The study has UK Research Ethics Committee approval (10/H0305/83, granted by the Cambridge South REC, and GEN/284/12 granted by the Republic of Ireland REC). The research team acknowledges the support of the National Institute for Health Research, through the Comprehensive Clinical Research Network. The Centers for Mendelian Genomics are funded by the National Human Genome Research Institute, the National Heart, Lung, and Blood Institute, and the National Eye Institute. Broad Institute (UM1 HG008900), Johns Hopkins University School of Medicine/Baylor College of Medicine (UM1 HG006542), University of Washington (UM1 HG006493), Yale University (UM1 HG006504).

## Contributions

P.C., V.MF, T.F.M, D.S. contributed to the data analysis, writing of the paper and design, execution of the work. S.A.M., M.E.D, M.B, K.A.P. contributed to data analysis and review of the manuscript. H.M., H.W., D.G.L, J.R.S contributed to data acquisition and data handling. T.K., H.H. contributed to development of the software and databases and review of the manuscript. J.P., V.N., T.S., C.-W.S., A.C, C.J.L, H.W-J., L.T., H.C, M.S, T.H. contributed to mouse phenotyping and data acquisition. H.F. led the mouse phenotyping. L.M.J.N. led the production of mouse models and cohorts for phenotyping and contributed to review of manuscript. A.M.F. led the mouse and embryo phenotyping and data acquisition and contributed to review of manuscript. V.G-D. led the mouse phenotyping and contributed to review of manuscript. D.S., T.F.M, A.L.B, J.D.H, M.H.d.A, W.W, Y.H, D.J.A, R.S, F.M, S.W, R.E.B, H.P., S.D.M.B., C.M., K.C.K.L. are PIs of the key programs that contributed to the management and execution of the work and contributed to review of the manuscript. The additional IMPC consortium members all contributed to data acquisition and data handling.

