## Supplementary Information for "Human and mouse essentiality screens as a resource for disease gene discovery"

#### Supplementary Tables

**Supplementary Table 1**. **Human cell essentiality assessment**. Comparison between the set of essential and non-essential genes based on mean Avana CRISPR-Cas9 screens performed on over 400 cell lines and 11 cell lines from 3 different studies (see Supplementary Fig. 1). For any given gene, a mean Avana score ≤ -0.45 resulted in considering the gene essential.

**Supplementary Table 2**. **Gene Ontology (GO) Biological Process (BP) Enrichment**. Set of Gene Ontology Biological Processes with significant results for the CL and DL category of genes (Fig. 1b). Genes in the SV and viable categories (VP, VN) were not significantly enriched in any biological process despite reasonable sample sizes, probably reflecting diverse roles for these genes.

**Supplementary Table 3**. **Embryo windows of lethality.** Embryonic viability assessment outcomes indicate the embryonic stage at which the homozygous LoF mice manifested lethality and their overlap with human cell essentiality categories. E, embryonic day.

**Supplementary Table 4. Gene features.** Adjusted P-values (Wilcoxon test, BH correction) for all pairwise comparisons (violin / boxplots in Fig. 2).

**Supplementary Table 5**. **Constraint scores**. Adjusted P-values (Wilcoxon test, BH correction) for all pairwise comparisons (violin / boxplots in Supplementary Fig. 2).

**Supplementary Table 6**. **Clinical features for AD disease genes across FUSIL bins.** Distribution of AD disease genes across FUSIL bins based on the number of physiological systems affected and the age of onset (only those genes with information for all three features were considered for this analysis). MoI, mode of inheritance; N, number of genes; Physiological systems affected: high (≥13), intermediate (6-13), low (≤6). Age of onset: early (antenatal, neonatal), intermediate (infancy, childhood), late (other).

**Supplementary Table 7. 163 DL genes that are highly intolerant to loss-of-function variation and not currently associated with human disease.** Set of prioritised genes with the corresponding intolerance to variation scores. Information about candidate variants present on these genes identified in any of the 3 sequencing programs investigated in the current study: Y, candidate variant identified / candidate variant absent from gnomAD; N, candidate variant present in gnomAD.

**Supplementary Table 8. Clinical description of patients with variants in *VPS4.*** Phenotypes reported for each patient, shared phenotypes in bold.

**Supplementary Table 9. Clinical description of patients with *de novo* variants in *TMEM63B*.** Phenotypes reported for each patient, shared phenotypes in bold.

**Supplementary Table 10**. **FUSIL categories.** Classification of genes based on the mouse embryo LoF viability and phenotypes as obtained by the IMPC and human cell essentiality scores (Avana) as obtained by the Broad Institute. In bold, categories shown in Table 1.

**Supplementary Table 11**. **IMPC and MGI viability assessment.** IMPC viability outcomes compared to MGI reported phenotypes.

**Supplementary Table 1**. **Human cell essentiality assessment**.

| Mean Avana -0.45 threshold | 11 cell lines | Number of overlapping genes | % Overlap | % total |
|---|---|---|---|---|
| Essential | Essential | 1,339 | 79.85 % | 96.11 % |
| Essential | Non-essential | 338 | 20.15 % | |
| Non-essential | Essential | 312 | 2.07 % | |
| Non-essential | Non-essential | 14,751 | 97.93 % | |

**Supplementary Table 2. Gene Ontology (GO) Biological Process (BP) Enrichment.**

| GOBPID | Term | Count | Size | Odds Ratio | P-value | P-value BH | FUSIL |
|---|---|---|---|---|---|---|---|
| GO:0090304 | nucleic acid metabolic process | 205 | 965 | 3.5 | 2.00E-29 | 6.30E-26 | CL |
| GO:0006807 | nitrogen compound metabolic process | 323 | 2134 | 2.8 | 2.30E-18 | 3.50E-15 | CL |
| GO:0000375 | RNA splicing, via transesterification reactions | 35 | 68 | 8.8 | 2.60E-16 | 2.70E-13 | CL |
| GO:0006403 | RNA localization | 28 | 49 | 11 | 7.10E-15 | 5.50E-12 | CL |
| GO:0006518 | peptide metabolic process | 54 | 159 | 4.4 | 1.20E-14 | 7.50E-12 | CL |
| GO:0043604 | amide biosynthetic process | 54 | 163 | 4.2 | 4.10E-14 | 1.80E-11 | CL |
| GO:0006396 | RNA processing | 31 | 68 | 7.9 | 4.10E-14 | 1.80E-11 | CL |
| GO:0044237 | cellular metabolic process | 58 | 306 | 4.5 | 9.30E-14 | 3.60E-11 | CL |
| GO:0071426 | ribonucleoprotein complex export from nucleus | 19 | 29 | 15.3 | 6.50E-12 | 2.20E-09 | CL |
| GO:0098781 | ncRNA transcription | 17 | 24 | 19.7 | 1.10E-11 | 3.40E-09 | CL |
| GO:0042795 | snRNA transcription by RNA polymerase II | 17 | 24 | 19.5 | 1.30E-11 | 3.60E-09 | CL |
| GO:0043039 | tRNA aminoacylation | 15 | 19 | 29.9 | 1.60E-11 | 4.20E-09 | CL |
| GO:0016073 | snRNA metabolic process | 14 | 19 | 22.8 | 2.90E-10 | 7.00E-08 | CL |
| GO:0015931 | nucleobase-containing compound transport | 23 | 50 | 6.9 | 6.90E-10 | 1.50E-07 | CL |
| GO:0051301 | cell division | 43 | 146 | 3.5 | 1.20E-09 | 2.50E-07 | CL |
| GO:0000398 | mRNA splicing, via spliceosome | 17 | 30 | 10.9 | 1.30E-09 | 2.50E-07 | CL |
| GO:0071840 | cellular component organization or biogenesis | 234 | 1550 | 1.9 | 2.40E-09 | 4.40E-07 | CL |
| GO:0051306 | mitotic sister chromatid separation | 14 | 22 | 13.9 | 7.70E-09 | 1.30E-06 | CL |
| GO:0006406 | mRNA export from nucleus | 13 | 20 | 14.8 | 1.80E-08 | 2.90E-06 | CL |
| GO:0070125 | mitochondrial translational elongation | 12 | 19 | 13.6 | 1.10E-07 | 1.60E-05 | CL |
| GO:0044770 | cell cycle phase transition | 24 | 69 | 4.5 | 1.10E-07 | 1.60E-05 | CL |
| GO:0006412 | translation | 11 | 17 | 15.9 | 1.20E-07 | 1.60E-05 | CL |
| GO:0044271 | cellular nitrogen compound biosynthetic process | 124 | 802 | 1.9 | 1.50E-07 | 2.00E-05 | CL |
| GO:0048285 | organelle fission | 34 | 119 | 3.3 | 1.90E-07 | 2.40E-05 | CL |
| GO:0006383 | transcription by RNA polymerase III | 7 | 7 | Inf | 2.30E-07 | 2.80E-05 | CL |
| GO:0006363 | termination of RNA polymerase I transcription | 8 | 9 | 62.8 | 2.30E-07 | 2.80E-05 | CL |
| GO:1901576 | organic substance biosynthetic process | 23 | 84 | 4.3 | 2.80E-07 | 3.20E-05 | CL |
| GO:0065004 | protein-DNA complex assembly | 16 | 35 | 6.8 | 3.30E-07 | 3.60E-05 | CL |
| GO:0007049 | cell cycle | 34 | 135 | 3.1 | 3.40E-07 | 3.60E-05 | CL |
| GO:0006281 | DNA repair | 26 | 83 | 3.7 | 6.70E-07 | 6.90E-05 | CL |
| GO:0006413 | translational initiation | 9 | 13 | 18.1 | 1.30E-06 | 1.30E-04 | CL |
| GO:0034660 | ncRNA metabolic process | 14 | 33 | 6.7 | 1.40E-06 | 1.40E-04 | CL |
| GO:0006283 | transcription-coupled nucleotide-excision repair | 9 | 13 | 17.7 | 1.50E-06 | 1.40E-04 | CL |
| GO:0019438 | aromatic compound biosynthetic process | 138 | 858 | 1.7 | 1.50E-06 | 1.40E-04 | CL |
| GO:0018130 | heterocycle biosynthetic process | 135 | 837 | 1.7 | 1.60E-06 | 1.40E-04 | CL |
| GO:0071826 | ribonucleoprotein complex subunit organization | 18 | 49 | 4.6 | 3.30E-06 | 2.90E-04 | CL |
| GO:0070126 | mitochondrial translational termination | 10 | 18 | 9.8 | 6.80E-06 | 5.70E-04 | CL |
| GO:1901362 | organic cyclic compound biosynthetic process | 137 | 874 | 1.7 | 7.00E-06 | 5.80E-04 | CL |
| GO:0006370 | 7-methylguanosine mRNA capping | 7 | 9 | 27.4 | 7.40E-06 | 5.80E-04 | CL |
| GO:0036260 | RNA capping | 7 | 9 | 27.4 | 7.40E-06 | 5.80E-04 | CL |

| | | | | | | | |
|---|---|---|---|---|---|---|---|
| GO:1901566 | organonitrogen compound biosynthetic process | 70 | 378 | 1.9 | 1.20E-05 | 9.10E-04 | CL |
| GO:0006361 | transcription initiation from RNA polymerase I promoter | 6 | 7 | 47.2 | 1.40E-05 | 1.00E-03 | CL |
| GO:0022613 | ribonucleoprotein complex biogenesis | 15 | 42 | 4.7 | 1.70E-05 | 1.20E-03 | CL |
| GO:0006888 | ER to Golgi vesicle-mediated transport | 16 | 46 | 4.2 | 2.50E-05 | 1.80E-03 | CL |
| GO:0008380 | RNA splicing | 7 | 11 | 14.8 | 3.50E-05 | 2.40E-03 | CL |
| GO:0031145 | anaphase-promoting complex-dependent catabolic process | 8 | 14 | 10.4 | 4.60E-05 | 3.00E-03 | CL |
| GO:0045841 | negative regulation of mitotic metaphase/anaphase transition | 8 | 14 | 10.4 | 4.60E-05 | 3.00E-03 | CL |
| GO:0071806 | protein transmembrane transport | 8 | 14 | 10.4 | 4.60E-05 | 3.00E-03 | CL |
| GO:0031440 | regulation of mRNA 3'-end processing | 6 | 8 | 23.6 | 4.90E-05 | 3.10E-03 | CL |
| GO:0006270 | DNA replication initiation | 6 | 8 | 23.4 | 5.10E-05 | 3.20E-03 | CL |
| GO:0019068 | virion assembly | 7 | 11 | 13.7 | 5.50E-05 | 3.40E-03 | CL |
| GO:0050658 | RNA transport | 8 | 15 | 9.3 | 6.80E-05 | 4.10E-03 | CL |
| GO:0006913 | nucleocytoplasmic transport | 11 | 27 | 5.6 | 7.50E-05 | 4.40E-03 | CL |
| GO:1901988 | negative regulation of cell cycle phase transition | 16 | 50 | 3.7 | 8.40E-05 | 4.90E-03 | CL |
| GO:0033048 | negative regulation of mitotic sister chromatid segregation | 8 | 15 | 8.9 | 8.90E-05 | 5.00E-03 | CL |
| GO:0050686 | negative regulation of mRNA processing | 5 | 6 | 39.1 | 1.00E-04 | 5.60E-03 | CL |
| GO:0000083 | regulation of transcription involved in G1/S transition of mitotic cell cycle | 5 | 6 | 38.9 | 1.10E-04 | 5.60E-03 | CL |
| GO:0032201 | telomere maintenance via semi-conservative replication | 5 | 6 | 38.9 | 1.10E-04 | 5.60E-03 | CL |
| GO:1900364 | negative regulation of mRNA polyadenylation | 5 | 6 | 38.9 | 1.10E-04 | 5.60E-03 | CL |
| GO:0009060 | aerobic respiration | 7 | 12 | 11 | 1.20E-04 | 6.10E-03 | CL |
| GO:0007059 | chromosome segregation | 16 | 53 | 3.6 | 1.20E-04 | 6.10E-03 | CL |
| GO:0000082 | G1/S transition of mitotic cell cycle | 9 | 20 | 6.6 | 1.50E-04 | 7.30E-03 | CL |
| GO:1905819 | negative regulation of chromosome separation | 8 | 16 | 7.8 | 1.60E-04 | 7.90E-03 | CL |
| GO:0043933 | protein-containing complex subunit organization | 64 | 393 | 1.8 | 1.70E-04 | 8.30E-03 | CL |
| GO:0033314 | mitotic DNA replication checkpoint | 4 | 4 | Inf | 1.70E-04 | 8.30E-03 | CL |
| GO:1903047 | mitotic cell cycle process | 20 | 82 | 2.8 | 2.20E-04 | 1.00E-02 | CL |
| GO:0071897 | DNA biosynthetic process | 16 | 55 | 3.2 | 3.00E-04 | 1.40E-02 | CL |
| GO:0010564 | regulation of cell cycle process | 25 | 111 | 2.4 | 3.10E-04 | 1.40E-02 | CL |
| GO:0009889 | regulation of biosynthetic process | 123 | 823 | 1.5 | 3.10E-04 | 1.40E-02 | CL |
| GO:0010972 | negative regulation of G2/M transition of mitotic cell cycle | 6 | 10 | 11.7 | 3.10E-04 | 1.40E-02 | CL |
| GO:0036258 | multivesicular body assembly | 6 | 10 | 11.7 | 3.10E-04 | 1.40E-02 | CL |
| GO:0051276 | chromosome organization | 9 | 24 | 5.4 | 3.40E-04 | 1.50E-02 | CL |
| GO:0042773 | ATP synthesis coupled electron transport | 10 | 26 | 4.9 | 3.50E-04 | 1.50E-02 | CL |
| GO:0022607 | cellular component assembly | 104 | 687 | 1.5 | 3.50E-04 | 1.50E-02 | CL |
| GO:0009057 | macromolecule catabolic process | 22 | 91 | 2.6 | 3.80E-04 | 1.60E-02 | CL |
| GO:0006368 | transcription elongation from RNA polymerase II promoter | 7 | 14 | 7.9 | 3.90E-04 | 1.60E-02 | CL |
| GO:0006890 | retrograde vesicle-mediated transport, Golgi to ER | 9 | 22 | 5.4 | 4.10E-04 | 1.60E-02 | CL |
| GO:0051783 | regulation of nuclear division | 11 | 32 | 4.2 | 4.90E-04 | 2.00E-02 | CL |
| GO:0071704 | organic substance metabolic process | 96 | 966 | 1.7 | 5.20E-04 | 2.10E-02 | CL |
| GO:0046700 | heterocycle catabolic process | 27 | 123 | 2.2 | 5.50E-04 | 2.10E-02 | CL |

| | | | | | | | |
|---|---|---|---|---|---|---|---|
| GO:0015980 | energy derivation by oxidation of organic compounds | 18 | 69 | 2.8 | 5.50E-04 | 2.10E-02 | CL |
| GO:0007080 | mitotic metaphase plate congression | 6 | 11 | 9.4 | 6.10E-04 | 2.30E-02 | CL |
| GO:0009126 | purine nucleoside monophosphate metabolic process | 18 | 70 | 2.7 | 6.70E-04 | 2.50E-02 | CL |
| GO:0009161 | ribonucleoside monophosphate metabolic process | 12 | 38 | 3.7 | 6.80E-04 | 2.50E-02 | CL |
| GO:0044270 | cellular nitrogen compound catabolic process | 27 | 125 | 2.2 | 7.20E-04 | 2.60E-02 | CL |
| GO:0033047 | regulation of mitotic sister chromatid segregation | 4 | 5 | 31.8 | 7.30E-04 | 2.60E-02 | CL |
| GO:1905818 | regulation of chromosome separation | 4 | 5 | 31.7 | 7.30E-04 | 2.60E-02 | CL |
| GO:0007094 | mitotic spindle assembly checkpoint | 4 | 5 | 31.3 | 7.60E-04 | 2.60E-02 | CL |
| GO:0031577 | spindle checkpoint | 4 | 5 | 31.3 | 7.60E-04 | 2.60E-02 | CL |
| GO:0000387 | spliceosomal snRNP assembly | 4 | 5 | 31.3 | 7.70E-04 | 2.60E-02 | CL |
| GO:0006297 | nucleotide-excision repair, DNA gap filling | 4 | 5 | 31.1 | 7.90E-04 | 2.60E-02 | CL |
| GO:0006369 | termination of RNA polymerase II transcription | 4 | 5 | 31.1 | 7.90E-04 | 2.60E-02 | CL |
| GO:0031442 | positive regulation of mRNA 3'-end processing | 4 | 5 | 31.1 | 7.90E-04 | 2.60E-02 | CL |
| GO:0080009 | mRNA methylation | 4 | 5 | 31.1 | 7.90E-04 | 2.60E-02 | CL |
| GO:0045292 | mRNA cis splicing, via spliceosome | 5 | 8 | 13 | 8.00E-04 | 2.60E-02 | CL |
| GO:0006296 | nucleotide-excision repair, DNA incision, 5'-to lesion | 5 | 8 | 13 | 8.20E-04 | 2.60E-02 | CL |
| GO:0032968 | positive regulation of transcription elongation from RNA polymerase II promoter | 5 | 8 | 13 | 8.20E-04 | 2.60E-02 | CL |
| GO:0019439 | aromatic compound catabolic process | 27 | 126 | 2.2 | 8.20E-04 | 2.60E-02 | CL |
| GO:0007093 | mitotic cell cycle checkpoint | 8 | 20 | 5.3 | 9.10E-04 | 2.90E-02 | CL |
| GO:0044419 | interspecies interaction between organisms | 31 | 155 | 2 | 1.00E-03 | 3.20E-02 | CL |
| GO:0006446 | regulation of translational initiation | 6 | 12 | 7.9 | 1.10E-03 | 3.30E-02 | CL |
| GO:0009144 | purine nucleoside triphosphate metabolic process | 12 | 40 | 3.4 | 1.20E-03 | 3.60E-02 | CL |
| GO:0009199 | ribonucleoside triphosphate metabolic process | 12 | 40 | 3.4 | 1.20E-03 | 3.60E-02 | CL |
| GO:0006139 | nucleobase-containing compound metabolic process | 21 | 146 | 2.3 | 1.20E-03 | 3.60E-02 | CL |
| GO:0006260 | DNA replication | 7 | 17 | 5.8 | 1.30E-03 | 3.90E-02 | CL |
| GO:0006399 | tRNA metabolic process | 3 | 3 | Inf | 1.30E-03 | 3.90E-02 | CL |
| GO:0010638 | positive regulation of organelle organization | 31 | 157 | 2 | 1.50E-03 | 3.90E-02 | CL |
| GO:0034472 | snRNA 3'-end processing | 3 | 3 | Inf | 1.50E-03 | 3.90E-02 | CL |
| GO:0001832 | blastocyst growth | 3 | 3 | Inf | 1.50E-03 | 3.90E-02 | CL |
| GO:0000244 | spliceosomal tri-snRNP complex assembly | 3 | 3 | Inf | 1.50E-03 | 3.90E-02 | CL |
| GO:0000463 | maturation of LSU-rRNA from tricistronic rRNA transcript (SSU-rRNA, 5.8S rRNA, LSU-rRNA) | 3 | 3 | Inf | 1.50E-03 | 3.90E-02 | CL |
| GO:0006265 | DNA topological change | 3 | 3 | Inf | 1.50E-03 | 3.90E-02 | CL |
| GO:0006999 | nuclear pore organization | 3 | 3 | Inf | 1.50E-03 | 3.90E-02 | CL |
| GO:0045039 | protein import into mitochondrial inner membrane | 3 | 3 | Inf | 1.50E-03 | 3.90E-02 | CL |
| GO:0071042 | nuclear polyadenylation-dependent mRNA catabolic process | 3 | 3 | Inf | 1.50E-03 | 3.90E-02 | CL |
| GO:0090166 | Golgi disassembly | 3 | 3 | Inf | 1.50E-03 | 3.90E-02 | CL |
| GO:0106074 | aminoacyl-tRNA metabolism involved in translational fidelity | 3 | 3 | Inf | 1.50E-03 | 3.90E-02 | CL |
| GO:1904263 | positive regulation of TORC1 signaling | 3 | 3 | Inf | 1.50E-03 | 3.90E-02 | CL |
| GO:1904874 | positive regulation of telomerase RNA localization to Cajal body | 3 | 3 | Inf | 1.50E-03 | 3.90E-02 | CL |

| | | | | | | | |
|---|---|---|---|---|---|---|---|
| GO:2001168 | positive regulation of histone H2B ubiquitination | 3 | 3 | Inf | 1.50E-03 | 3.90E-02 | CL |
| GO:0006323 | DNA packaging | 5 | 9 | 9.9 | 1.60E-03 | 4.00E-02 | CL |
| GO:1901361 | organic cyclic compound catabolic process | 27 | 132 | 2 | 1.70E-03 | 4.40E-02 | CL |
| GO:0043044 | ATP-dependent chromatin remodeling | 7 | 17 | 5.5 | 1.80E-03 | 4.40E-02 | CL |
| GO:0042307 | positive regulation of protein import into nucleus | 6 | 13 | 6.7 | 1.90E-03 | 4.70E-02 | CL |
| GO:0009792 | embryo development ending in birth or egg hatching | 72 | 127 | 5.4 | 2.60E-19 | 1.40E-15 | DL |
| GO:0001701 | in utero embryonic development | 28 | 41 | 8.5 | 5.50E-11 | 1.50E-07 | DL |
| GO:0048736 | appendage development | 25 | 35 | 9.7 | 1.70E-10 | 3.00E-07 | DL |
| GO:0034645 | cellular macromolecule biosynthetic process | 265 | 970 | 1.6 | 1.80E-08 | 2.40E-05 | DL |
| GO:0051254 | positive regulation of RNA metabolic process | 117 | 356 | 2 | 2.30E-08 | 2.50E-05 | DL |
| GO:0010557 | positive regulation of macromolecule biosynthetic process | 121 | 375 | 2 | 3.40E-08 | 3.00E-05 | DL |
| GO:0006996 | organelle organization | 101 | 317 | 2.1 | 4.70E-08 | 3.40E-05 | DL |
| GO:0048729 | tissue morphogenesis | 50 | 118 | 2.9 | 5.10E-08 | 3.40E-05 | DL |
| GO:0031328 | positive regulation of cellular biosynthetic process | 124 | 397 | 1.9 | 1.90E-07 | 1.10E-04 | DL |
| GO:0072175 | epithelial tube formation | 17 | 25 | 8.2 | 4.70E-07 | 2.50E-04 | DL |
| GO:0001841 | neural tube formation | 16 | 23 | 8.8 | 6.90E-07 | 3.20E-04 | DL |
| GO:0016331 | morphogenesis of embryonic epithelium | 18 | 28 | 7 | 7.30E-07 | 3.20E-04 | DL |
| GO:0034654 | nucleobase-containing compound biosynthetic process | 226 | 844 | 1.5 | 2.40E-06 | 9.70E-04 | DL |
| GO:0042733 | embryonic digit morphogenesis | 8 | 8 | Inf | 3.60E-06 | 1.40E-03 | DL |
| GO:0045944 | positive regulation of transcription by RNA polymerase II | 76 | 230 | 2 | 6.20E-06 | 2.20E-03 | DL |
| GO:0009799 | specification of symmetry | 19 | 35 | 4.6 | 1.40E-05 | 4.50E-03 | DL |
| GO:0009058 | biosynthetic process | 121 | 453 | 1.7 | 1.40E-05 | 4.50E-03 | DL |
| GO:0021904 | dorsal/ventral neural tube patterning | 7 | 7 | Inf | 1.80E-05 | 5.20E-03 | DL |
| GO:0032981 | mitochondrial respiratory chain complex I assembly | 11 | 15 | 10.5 | 2.00E-05 | 5.20E-03 | DL |
| GO:1902679 | negative regulation of RNA biosynthetic process | 81 | 255 | 1.8 | 2.00E-05 | 5.20E-03 | DL |
| GO:0042475 | odontogenesis of dentin-containing tooth | 14 | 23 | 6 | 3.50E-05 | 8.10E-03 | DL |
| GO:0001843 | neural tube closure | 12 | 18 | 7.7 | 3.50E-05 | 8.10E-03 | DL |
| GO:1903363 | negative regulation of cellular protein catabolic process | 14 | 23 | 5.9 | 3.50E-05 | 8.10E-03 | DL |
| GO:0150063 | visual system development | 32 | 78 | 2.7 | 4.00E-05 | 8.90E-03 | DL |
| GO:0060976 | coronary vasculature development | 11 | 16 | 8.4 | 5.10E-05 | 1.10E-02 | DL |
| GO:0000122 | negative regulation of transcription by RNA polymerase II | 60 | 181 | 2 | 5.50E-05 | 1.10E-02 | DL |
| GO:0045934 | negative regulation of nucleobase-containing compound metabolic process | 90 | 300 | 1.7 | 7.90E-05 | 1.60E-02 | DL |
| GO:0021983 | pituitary gland development | 9 | 12 | 11.4 | 9.10E-05 | 1.70E-02 | DL |
| GO:0048538 | thymus development | 9 | 12 | 11.4 | 9.10E-05 | 1.70E-02 | DL |
| GO:0035116 | embryonic hindlimb morphogenesis | 8 | 10 | 15.2 | 1.10E-04 | 1.90E-02 | DL |
| GO:1904837 | beta-catenin-TCF complex assembly | 8 | 10 | 15.2 | 1.10E-04 | 1.90E-02 | DL |
| GO:0060325 | face morphogenesis | 7 | 8 | 26.5 | 1.20E-04 | 1.90E-02 | DL |
| GO:0007368 | determination of left/right symmetry | 15 | 28 | 4.4 | 1.40E-04 | 2.20E-02 | DL |
| GO:0072163 | mesonephric epithelium development | 12 | 20 | 5.7 | 1.60E-04 | 2.40E-02 | DL |
| GO:0009108 | coenzyme biosynthetic process | 10 | 15 | 7.7 | 1.60E-04 | 2.40E-02 | DL |

| | | | | | | | |
|---|---|---|---|---|---|---|---|
| GO:0061326 | renal tubule development | 13 | 23 | 5 | 2.00E-04 | 2.90E-02 | DL |
| GO:0018394 | peptidyl-lysine acetylation | 18 | 38 | 3.4 | 2.40E-04 | 3.40E-02 | DL |
| GO:0042157 | lipoprotein metabolic process | 16 | 32 | 3.8 | 2.50E-04 | 3.40E-02 | DL |
| GO:0006475 | internal protein amino acid acetylation | 19 | 42 | 3.2 | 3.40E-04 | 4.70E-02 | DL |
| GO:0001570 | vasculogenesis | 10 | 16 | 6.4 | 3.60E-04 | 4.70E-02 | DL |

**Supplementary Table 3**. **Embryo windows of lethality**.

<table>
<tr><th rowspan="2">Mouse embryonic group</th><th rowspan="2">Windows of embryo lethality</th><th rowspan="2">Total number of genes</th><th rowspan="2">%</th><th colspan="2">Genes with human cell essentiality information (%)</th></tr>
<tr><th>Essential</th><th>Non-essential</th></tr>
<tr><td>Early gestation</td><td>prior to E9.5</td><td>197</td><td>49.25%</td><td>125 (64.76%)</td><td>68 (35.23%)</td></tr>
<tr><td rowspan="2">Mid gestation</td><td>E9.5-E12.5</td><td>45</td><td rowspan="2">12.50%</td><td rowspan="2">5 (10.20%)</td><td rowspan="2">44 (89.80%)</td></tr>
<tr><td>E12.5-E14.5/E15.5</td><td>5</td></tr>
<tr><td rowspan="3">Late gestation</td><td>E14.5/E15.5-E18.5</td><td>3</td><td rowspan="3">38.25%</td><td rowspan="3">7 (4.70%)</td><td rowspan="3">142 (95.30%)</td></tr>
<tr><td>after E14.5/E15.5</td><td>75</td></tr>
<tr><td>after E18.5</td><td>75</td></tr>
</table>

**Supplementary Table 4. Gene features.**

| FUSIL bin 1 | FUSIL bin 2 | Recomb Rate | TPM Brain Cortex | TPM Cells Transform Fibroblasts | TPM Ovary | TPM Testis | Degree | Topological Coefficient | Probability of mutation | Transcript length | GIMS Selection Score | pLI |
|---|---|---|---|---|---|---|---|---|---|---|---|---|
| CL | DL | 5.7E-16 | 1.8E-06 | 3.3E-15 | 4.9E-09 | 5.5E-19 | 1.4E-16 | 3.1E-03 | 4.0E-01 | 3.3E-02 | 7.8E-01 | 1.1E-01 |
| CL | SV | 5.3E-15 | 2.1E-10 | 8.0E-21 | 3.3E-14 | 1.4E-20 | 2.0E-17 | 6.7E-06 | 6.0E-02 | 1.2E-04 | 7.8E-01 | 1.6E-01 |
| CL | VP | 1.0E-33 | 6.4E-30 | 3.4E-77 | 4.1E-51 | 2.5E-57 | 5.2E-46 | 2.3E-12 | 1.7E-01 | 3.4E-01 | 5.6E-18 | 9.7E-12 |
| CL | VN | 8.0E-21 | 8.0E-22 | 1.0E-51 | 9.0E-38 | 9.9E-39 | 2.2E-24 | 3.9E-10 | 3.5E-02 | 7.7E-02 | 5.0E-12 | 2.2E-08 |
| DL | SV | 4.8E-01 | 2.0E-02 | 1.0E-03 | 2.6E-03 | 1.7E-02 | 1.3E-02 | 4.0E-02 | 1.3E-01 | 3.2E-02 | 7.8E-01 | 9.7E-01 |
| DL | VP | 2.3E-04 | 3.4E-16 | 6.9E-43 | 1.1E-31 | 1.4E-20 | 5.0E-16 | 6.7E-06 | 4.8E-03 | 4.4E-05 | 1.2E-30 | 5.3E-28 |
| DL | VN | 1.0E-02 | 1.2E-11 | 4.9E-28 | 1.7E-22 | 4.5E-15 | 3.4E-08 | 4.9E-05 | 1.4E-03 | 1.1E-04 | 1.0E-15 | 2.7E-15 |
| SV | VP | 4.0E-02 | 2.2E-05 | 1.1E-12 | 2.9E-09 | 4.3E-06 | 2.6E-04 | 2.3E-01 | 3.9E-05 | 2.8E-08 | 1.2E-20 | 1.6E-17 |
| SV | VN | 8.8E-02 | 7.5E-06 | 5.3E-13 | 3.2E-10 | 7.9E-07 | 1.7E-03 | 6.4E-02 | 2.8E-05 | 2.0E-07 | 2.2E-13 | 4.9E-12 |
| VP | VN | 9.1E-01 | 1.0E-01 | 6.2E-03 | 1.3E-02 | 5.1E-02 | 4.9E-01 | 2.4E-01 | 1.3E-01 | 1.9E-01 | 5.7E-01 | 7.2E-01 |

**Supplementary Table 5**. **Constraint scores**.

| group1 | group2 | o/e LoF | o/e LoF upper bound (LOEUF) | o/e mis | o/e syn | shet | RVIS | HI |
|---|---|---|---|---|---|---|---|---|
| CL | DL | 4.1E-01 | 5.0E-01 | 5.6E-01 | 9.6E-01 | 8.9E-01 | 1.4E-01 | 3.7E-01 |
| CL | SV | 6.8E-01 | 3.6E-01 | 5.6E-01 | 5.2E-01 | 5.8E-01 | 7.9E-01 | 1.2E-01 |
| CL | VP | 3.4E-21 | 4.7E-23 | 3.3E-14 | 2.1E-01 | 8.1E-19 | 1.8E-20 | 9.2E-42 |
| CL | VN | 1.2E-16 | 2.3E-21 | 2.0E-11 | 5.2E-01 | 1.8E-15 | 6.7E-18 | 2.8E-28 |
| DL | SV | 6.8E-01 | 7.3E-01 | 9.1E-01 | 5.2E-01 | 5.6E-01 | 9.8E-02 | 3.7E-01 |
| DL | VP | 1.1E-38 | 2.0E-38 | 5.8E-20 | 1.3E-01 | 9.4E-26 | 1.8E-20 | 1.3E-52 |
| DL | VN | 4.6E-23 | 2.7E-27 | 4.0E-13 | 5.2E-01 | 2.1E-17 | 1.6E-15 | 1.2E-28 |
| SV | VP | 1.8E-22 | 3.0E-26 | 3.3E-13 | 5.2E-01 | 7.7E-21 | 9.6E-20 | 3.6E-29 |
| SV | VN | 2.0E-17 | 3.8E-23 | 7.0E-11 | 9.6E-01 | 2.1E-17 | 1.1E-16 | 6.3E-21 |
| VP | VN | 1.3E-01 | 3.8E-03 | 2.0E-01 | 5.2E-01 | 5.3E-02 | 7.5E-02 | 1.2E-01 |

**Supplementary Table 6**. **Clinical features for AD disease genes across FUSIL bins.**

| FUSIL | MoI | Physiological systems affected | Age of onset | N | N FUSIL | N FUSIL and MoI | % (FUSIL) | % (FUSIL and MoI) |
|---|---|---|---|---|---|---|---|---|
| CL | AD | high | early | 5 | 110 | 22 | 4.55 | 22.73 |
| CL | AD | high | intermediate | 2 | 110 | 22 | 1.82 | 9.09 |
| CL | AD | high | late | 0 | 110 | 22 | 0 | 0 |
| CL | AD | intermediate | early | 2 | 110 | 22 | 1.82 | 9.09 |
| CL | AD | intermediate | intermediate | 5 | 110 | 22 | 4.55 | 22.73 |
| CL | AD | intermediate | late | 3 | 110 | 22 | 2.73 | 13.64 |
| CL | AD | low | early | 1 | 110 | 22 | 0.91 | 4.55 |
| CL | AD | low | intermediate | 3 | 110 | 22 | 2.73 | 13.64 |
| CL | AD | low | late | 1 | 110 | 22 | 0.91 | 4.55 |
| DL | AD | high | early | 24 | 264 | 82 | 9.09 | 29.27 |
| DL | AD | high | intermediate | 6 | 264 | 82 | 2.27 | 7.32 |
| DL | AD | high | late | 2 | 264 | 82 | 0.76 | 2.44 |
| DL | AD | intermediate | early | 16 | 264 | 82 | 6.06 | 19.51 |
| DL | AD | intermediate | intermediate | 6 | 264 | 82 | 2.27 | 7.32 |
| DL | AD | intermediate | late | 6 | 264 | 82 | 2.27 | 7.32 |
| DL | AD | low | early | 9 | 264 | 82 | 3.41 | 10.98 |
| DL | AD | low | intermediate | 9 | 264 | 82 | 3.41 | 10.98 |
| DL | AD | low | late | 4 | 264 | 82 | 1.52 | 4.88 |
| SV | AD | high | early | 8 | 113 | 22 | 7.08 | 36.36 |
| SV | AD | high | intermediate | 1 | 113 | 22 | 0.88 | 4.55 |
| SV | AD | high | late | 0 | 113 | 22 | 0 | 0 |
| SV | AD | intermediate | early | 6 | 113 | 22 | 5.31 | 27.27 |
| SV | AD | intermediate | intermediate | 1 | 113 | 22 | 0.88 | 4.55 |
| SV | AD | intermediate | late | 1 | 113 | 22 | 0.88 | 4.55 |
| SV | AD | low | early | 2 | 113 | 22 | 1.77 | 9.09 |
| SV | AD | low | intermediate | 1 | 113 | 22 | 0.88 | 4.55 |
| SV | AD | low | late | 2 | 113 | 22 | 1.77 | 9.09 |
| VP | AD | high | early | 8 | 288 | 70 | 2.78 | 11.43 |
| VP | AD | high | intermediate | 3 | 288 | 70 | 1.04 | 4.29 |
| VP | AD | high | late | 4 | 288 | 70 | 1.39 | 5.71 |
| VP | AD | intermediate | early | 12 | 288 | 70 | 4.17 | 17.14 |
| VP | AD | intermediate | intermediate | 5 | 288 | 70 | 1.74 | 7.14 |
| VP | AD | intermediate | late | 9 | 288 | 70 | 3.12 | 12.86 |
| VP | AD | low | early | 6 | 288 | 70 | 2.08 | 8.57 |
| VP | AD | low | intermediate | 8 | 288 | 70 | 2.78 | 11.43 |
| VP | AD | low | late | 15 | 288 | 70 | 5.21 | 21.43 |
| VN | AD | high | early | 0 | 28 | 14 | 0 | 0 |
| VN | AD | high | intermediate | 0 | 28 | 14 | 0 | 0 |
| VN | AD | high | late | 0 | 28 | 14 | 0 | 0 |
| VN | AD | intermediate | early | 2 | 28 | 14 | 7.14 | 14.29 |
| VN | AD | intermediate | intermediate | 3 | 28 | 14 | 10.71 | 21.43 |
| VN | AD | intermediate | late | 1 | 28 | 14 | 3.57 | 7.14 |
| VN | AD | low | early | 1 | 28 | 14 | 3.57 | 7.14 |
| VN | AD | low | intermediate | 3 | 28 | 14 | 10.71 | 21.43 |
| VN | AD | low | late | 4 | 28 | 14 | 14.29 | 28.57 |

**Supplementary Table 7. 163 DL genes that are highly intolerant to loss-of-function variation and not currently associated with human disease.**

| HGNC ID | Gene symbol | HI (<10) | pLI (> 0.90) | o/e LoF upper bound (<0.35) | o/e LoF | o/e mis | 100KGP candidates | 100KGP variants absent from gnomAD | DDD candidates | DDD variants absent from gnomAD | CMG candidates |
|---|---|---|---|---|---|---|---|---|---|---|---|
| HGNC:13488 | *VPS4A* | 17.58 | 0.928 | 0.36 | 0.139 | 0.532 | Y | Y | | | Y |
| HGNC:17735 | *TMEM63B* | 34.88 | 1 | 0.173 | 0.0668 | 0.475 | Y | Y | Y | Y | |
| HGNC:865 | *ATP6V0A1* | 20.07 | 0.998 | 0.27 | 0.144 | 0.517 | Y | Y | Y | Y | |
| HGNC:13731 | *MAEA* | 7.13 | 0.636 | 0.444 | 0.194 | 0.55 | Y | Y | Y | Y | |
| HGNC:24319 | *CMIP* | 19.68 | 1 | 0.149 | 0.0473 | 0.554 | Y | Y | Y | Y | |
| HGNC:9406 | *PKN2* | 12.8 | 1 | 0.154 | 0.0595 | 0.621 | Y | Y | Y | Y | |
| HGNC:11275 | *SPTBN1* | 2.63 | 1 | 0.077 | 0.0337 | 0.654 | Y | Y | Y | Y | |
| HGNC:9907 | *RBMS1* | 4.97 | 0.147 | 0.476 | 0.253 | 0.685 | Y | Y | Y | Y | |
| HGNC:14958 | *PUM2* | 8.73 | 0.994 | 0.286 | 0.164 | 0.688 | Y | Y | Y | Y | |
| HGNC:15909 | *NCOA5* | 21.03 | 0.999 | 0.214 | 0.068 | 0.851 | Y | Y | | | Y |
| HGNC:15985 | *SVEP1* | 50.43 | 1 | 0.248 | 0.181 | 0.862 | Y | Y | Y | Y | Y |
| HGNC:17798 | *YLPM1* | 23.71 | 1 | 0.233 | 0.151 | 0.925 | Y | Y | Y | Y | |
| HGNC:1779 | *CDK8* | 1.81 | 0.383 | 0.423 | 0.225 | 0.34 | Y | Y | Y | N | |
| HGNC:555 | *AP1G1* | 9.26 | 1 | 0.174 | 0.076 | 0.594 | Y | Y | Y | N | |
| HGNC:24659 | *EXOC8* | 44.12 | 0.998 | 0.206 | 0.0435 | 0.713 | Y | Y | Y | N | |
| HGNC:2556 | *CUL5* | 9.27 | 1 | 0.159 | 0.0615 | 0.33 | Y | Y | | | |
| HGNC:23393 | *CARM1* | 14.83 | 1 | 0.1 | 0 | 0.378 | Y | Y | | | |
| HGNC:29509 | *APH1A* | 9.08 | 0.75 | 0.476 | 0.151 | 0.548 | Y | Y | | | |
| HGNC:25416 | *ATXN7L3* | 34.92 | 0.999 | 0.193 | 0.0407 | 0.608 | Y | Y | | | |
| HGNC:23572 | *MORC3* | 33.9 | 1 | 0.194 | 0.0846 | 0.627 | Y | Y | | | |
| HGNC:24511 | *CHTOP* | 7.59 | 0.708 | 0.454 | 0.176 | 0.629 | Y | Y | | | |
| HGNC:6851 | *MAP3K12* | 24.82 | 1 | 0.073 | 0 | 0.641 | Y | Y | | | |
| HGNC:17620 | *NDEL1* | 11.34 | 0.98 | 0.304 | 0.0967 | 0.654 | Y | Y | | | |
| HGNC:10717 | *SEL1L* | 18.54 | 0.997 | 0.274 | 0.146 | 0.654 | Y | Y | | | |
| HGNC:19143 | *USP32* | 18.55 | 1 | 0.175 | 0.1 | 0.663 | Y | Y | | | |
| HGNC:9681 | *PTPRS* | 66.22 | 1 | 0.217 | 0.134 | 0.695 | Y | Y | | | |
| HGNC:2734 | *DDX1* | 8.05 | 0.999 | 0.252 | 0.128 | 0.726 | Y | Y | | | |
| HGNC:24224 | *CDK12* | 16.18 | 1 | 0.127 | 0.0493 | 0.737 | Y | Y | | | |
| HGNC:25067 | *INTS12* | 19.51 | 0.938 | 0.358 | 0.114 | 0.744 | Y | Y | | | |
| HGNC:9618 | *PTK7* | 29.42 | 0.997 | 0.275 | 0.152 | 0.759 | Y | Y | | | |
| HGNC:23035 | *L3MBTL3* | 25.7 | 1 | 0.239 | 0.114 | 0.769 | Y | Y | | | |
| HGNC:28483 | *TMEM161B* | 9.65 | 0.0000739 | 0.672 | 0.414 | 0.779 | Y | Y | | | |
| HGNC:4163 | *GART* | 9.36 | 0.000297 | 0.503 | 0.327 | 0.849 | Y | Y | | | |
| HGNC:20445 | *MBD6* | 33.61 | 1 | 0.205 | 0.0652 | 1.08 | Y | Y | | | |
| HGNC:6485 | *LAMA5* | 63.61 | 0.998 | 0.269 | 0.204 | 0.98 | Y | N | Y | N | Y |
| HGNC:25548 | *INTS10* | 35.08 | 1 | 0.164 | 0.0521 | 0.791 | Y | N | Y | Y | |

| | | | | | | | | | | | |
|---|---|---|---|---|---|---|---|---|---|---|---|
| HGNC:31104 | *MYO18A* | 15.46 | 0.976 | 0.292 | 0.201 | 0.793 | Y | N | Y | Y | |
| HGNC:12480 | *UBE2F* | 38.69 | 0.933 | 0.367 | 0.0773 | 0.631 | Y | N | Y | N | |
| HGNC:2904 | *DLG5* | 39.42 | 1 | 0.253 | 0.159 | 0.779 | Y | N | Y | N | |
| HGNC:172 | *ACVR1B* | 11.26 | 0.999 | 0.187 | 0.0395 | 0.464 | Y | N | | | |
| HGNC:9847 | *RANBP1* | 8.76 | 0.834 | 0.465 | 0.098 | 0.486 | Y | N | | | |
| HGNC:6847 | *MAP2K7* | 33.11 | 0.999 | 0.148 | 0 | 0.534 | Y | N | | | |
| HGNC:23040 | *BEND3* | 34.49 | 0.936 | 0.348 | 0.152 | 0.725 | Y | N | | | |
| HGNC:26002 | *CASZ1* | 12.36 | 1 | 0.15 | 0.0654 | 0.765 | Y | N | | | |
| HGNC:6163 | *ITGB8* | 16.77 | 0.999 | 0.233 | 0.102 | 0.86 | Y | N | | | |
| HGNC:25484 | *KIF26B* | 31.3 | 1 | 0.214 | 0.114 | 0.899 | Y | N | | | |
| HGNC:7777 | *NFATC3* | 7.9 | 0.998 | 0.267 | 0.135 | 0.938 | Y | N | | | |
| HGNC:28121 | *PRRC2B* | 43.15 | 1 | 0.186 | 0.117 | 0.941 | | | Y | Y | Y |
| HGNC:4216 | *GDF11* | 14.75 | 0.98 | 0.291 | 0.0614 | 0.444 | | | | | Y |
| HGNC:7880 | *CNOT4* | 6.19 | 1 | 0.14 | 0.0296 | 0.5 | | | | | Y |
| HGNC:11491 | *SYK* | 21.47 | 1 | 0.185 | 0.0587 | 0.583 | | | | | Y |
| HGNC:19355 | *JMJD6* | 24.76 | 0.933 | 0.357 | 0.138 | 0.732 | | | | | Y |
| HGNC:29093 | *ZC3H11A* | 39.28 | 1 | 0.184 | 0.0711 | 0.76 | | | | | Y |
| HGNC:11279 | *SQLE* | 42.86 | 0.97 | 0.324 | 0.125 | 0.798 | | | | | Y |
| HGNC:16051 | *PARD3* | 7.68 | 0.936 | 0.311 | 0.199 | 0.885 | | | | | Y |
| HGNC:17808 | *ZC3H4* | 56.54 | 1 | 0.056 | 0 | 0.931 | | | | | Y |
| HGNC:20271 | *NGDN* | 5.46 | 2.41E-08 | 1.070 | 0.694 | 1.01 | | | | | Y |
| HGNC:11254 | *SPOP* | 4.35 | 0.999 | 0.141 | 0 | 0.211 | | | Y | Y | |
| HGNC:4037 | *FYN* | 0.92 | 0.995 | 0.275 | 0.12 | 0.355 | | | Y | Y | |
| HGNC:563 | *AP2B1* | 6.87 | 0.998 | 0.267 | 0.135 | 0.367 | | | Y | Y | |
| HGNC:4395 | *GNAZ* | 36.02 | 0.963 | 0.294 | 0 | 0.406 | | | Y | Y | |
| HGNC:30092 | *NAMPT* | 8.84 | 0.982 | 0.305 | 0.118 | 0.458 | | | Y | Y | |
| HGNC:8634 | *PBX3* | 0.89 | 0.378 | 0.474 | 0.225 | 0.602 | | | Y | Y | |
| HGNC:24712 | *TENT5C* | 35.66 | 0.949 | 0.321 | 0 | 0.63 | | | Y | Y | |
| HGNC:12612 | *USP14* | 4.57 | 0.917 | 0.345 | 0.175 | 0.647 | | | Y | Y | |
| HGNC:4568 | *RAPGEF1* | 23.82 | 1 | 0.223 | 0.119 | 0.649 | | | Y | Y | |
| HGNC:25555 | *RIC8B* | 7.79 | 0.998 | 0.209 | 0.0441 | 0.651 | | | Y | Y | |
| HGNC:2663 | *DACH1* | 0.75 | 1 | 0.136 | 0 | 0.66 | | | Y | Y | |
| HGNC:7985 | *NR6A1* | 5.06 | 0.98 | 0.309 | 0.119 | 0.664 | | | Y | Y | |
| HGNC:8568 | *FURIN* | 5.18 | 1 | 0.192 | 0.0744 | 0.686 | | | Y | Y | |
| HGNC:6189 | *JAG2* | 35.89 | 1 | 0.137 | 0.0599 | 0.738 | | | Y | Y | |
| HGNC:18142 | *DAAM1* | 11.09 | 0.999 | 0.262 | 0.15 | 0.773 | | | Y | Y | |
| HGNC:27223 | *RBM33* | 42.71 | 1 | 0.051 | 0 | 0.778 | | | Y | Y | |
| HGNC:8004 | *NRP1* | 2.53 | 0.997 | 0.275 | 0.146 | 0.807 | | | Y | Y | |
| HGNC:18118 | *RSF1* | 34.22 | 1 | 0.043 | 0 | 0.808 | | | Y | Y | |
| HGNC:30291 | *G3BP2* | 6.8 | 1 | 0.177 | 0.0372 | 0.392 | | | Y | N | |
| HGNC:11241 | *SPI1* | 18.35 | 0.983 | 0.243 | 0 | 0.474 | | | Y | N | |
| HGNC:19678 | *UCHL5* | 5.6 | 0.00516 | 0.636 | 0.353 | 0.569 | | | Y | N | |

| | | | | | | | | | | |
|---|---|---|---|---|---|---|---|---|---|---|
| HGNC:11467 | *SUPT4H1* | 5.82 | 0.000617 | 1.320 | 0.668 | 0.603 | | | Y | N |
| HGNC:557 | *SYNRG* | 26.86 | 0.932 | 0.320 | 0.193 | 0.831 | | | Y | N |
| HGNC:1103 | *BRD2* | 1.63 | 1 | 0.217 | 0.0841 | 0.935 | | | Y | N |
| HGNC:11466 | *SUPT3H* | 4.33 | 2.9E-10 | 1.210 | 0.806 | 0.975 | | | Y | N |
| HGNC:9763 | *RAB2A* | 11.01 | 0.989 | 0.220 | 0 | 0.2 | | | | |
| HGNC:3312 | *ELAVL1* | 0.91 | 0.989 | 0.223 | 0 | 0.275 | | | | |
| HGNC:1551 | *CBX1* | 4.23 | 0.978 | 0.257 | 0 | 0.283 | | | | |
| HGNC:20995 | *ASF1A* | 1.71 | 0.758 | 0.530 | 0.112 | 0.329 | | | | |
| HGNC:2444 | *CSK* | 20.75 | 1 | 0.166 | 0.0351 | 0.34 | | | | |
| HGNC:7000 | *MEIS1* | 0.58 | 0.999 | 0.148 | 0 | 0.37 | | | | |
| HGNC:9304 | *PPP2R2A* | 4.71 | 0.995 | 0.255 | 0.081 | 0.385 | | | | |
| HGNC:14014 | *MEMO1* | 4 | 0.994 | 0.244 | 0.0514 | 0.401 | | | | |
| HGNC:690 | *ARIH2* | 17.16 | 0.996 | 0.276 | 0.131 | 0.406 | | | | |
| HGNC:2422 | *CS* | 7.67 | 0.983 | 0.306 | 0.134 | 0.407 | | | | |
| HGNC:24824 | *FZR1* | 42.53 | 1 | 0.169 | 0.0355 | 0.418 | | | | |
| HGNC:9774 | *RAB35* | 29.36 | 0.979 | 0.253 | 0 | 0.421 | | | | |
| HGNC:24590 | *DPY30* | 4.94 | 0.192 | 0.954 | 0.303 | 0.429 | | | | |
| HGNC:4071 | *GABPA* | 16.15 | 0.998 | 0.235 | 0.0747 | 0.433 | | | | |
| HGNC:705 | *ARPC2* | 6.72 | 0.998 | 0.167 | 0 | 0.441 | | | | |
| HGNC:29119 | *UBXN7* | 16.72 | 1 | 0.118 | 0 | 0.45 | | | | |
| HGNC:13724 | *ATP6V0D1* | 21.73 | 0.981 | 0.290 | 0.0611 | 0.458 | | | | |
| HGNC:14015 | *CELF4* | 4.37 | 0.997 | 0.246 | 0.0781 | 0.481 | | | | |
| HGNC:4852 | *HDAC1* | 0.32 | 0.537 | 0.418 | 0.212 | 0.505 | | | | |
| HGNC:12507 | *UBP1* | 26.92 | 1 | 0.086 | 0 | 0.505 | | | | |
| HGNC:23303 | *TSPAN14* | 31.5 | 0.975 | 0.303 | 0.0638 | 0.519 | | | | |
| HGNC:17615 | *PCGF1* | 24.56 | 0.997 | 0.172 | 0 | 0.524 | | | | |
| HGNC:10473 | *RUNX3* | 9.86 | 0.964 | 0.323 | 0.0681 | 0.524 | | | | |
| HGNC:6350 | *KLF7* | 5.04 | 0.98 | 0.253 | 0 | 0.529 | | | | |
| HGNC:23077 | *OTUB1* | 48.05 | 0.968 | 0.317 | 0.0668 | 0.531 | | | | |
| HGNC:18752 | *MAPKAP1* | 1.7 | 1 | 0.096 | 0 | 0.538 | | | | |
| HGNC:11108 | *SMARCD3* | 7.16 | 0.407 | 0.467 | 0.222 | 0.54 | | | | |
| HGNC:30231 | *TIPRL* | 25.12 | 0.942 | 0.356 | 0.0751 | 0.557 | | | | |
| HGNC:17346 | *PRPF4B* | 3.38 | 1 | 0.118 | 0.0376 | 0.562 | | | | |
| HGNC:13071 | *PATZ1* | 31.32 | 0.995 | 0.238 | 0.0503 | 0.572 | | | | |
| HGNC:32669 | *TRIM71* | 36.31 | 1 | 0.177 | 0.0373 | 0.573 | | | | |
| HGNC:744 | *ASH2L* | 19.21 | 1 | 0.211 | 0.0818 | 0.577 | | | | |
| HGNC:28412 | *NABP2* | 41.31 | 0.972 | 0.274 | 0 | 0.581 | | | | |
| HGNC:15630 | *TCERG1* | 3.91 | 1 | 0.223 | 0.124 | 0.59 | | | | |
| HGNC:17943 | *DERL2* | 12.71 | 0.956 | 0.336 | 0.0708 | 0.601 | | | | |
| HGNC:24543 | *EPC2* | 6.63 | 1 | 0.160 | 0.0507 | 0.611 | | | | |
| HGNC:20161 | *TOX4* | 22.69 | 0.964 | 0.328 | 0.143 | 0.621 | | | | |
| HGNC:15750 | *DSTN* | 8.92 | 0.557 | 0.722 | 0.152 | 0.622 | | | | |

| | | | | | | |
|---|---|---|---|---|---|---|
| HGNC:21168 | *RHOT1* | 7.79 | 0.762 | 0.355 | 0.203 | 0.626 |
| HGNC:13291 | *GABARAPL2* | 9.32 | 0.00023 | 1.590 | 0.829 | 0.63 |
| HGNC:13058 | *ZRANB2* | 5.99 | 1 | 0.181 | 0.0382 | 0.656 |
| HGNC:15607 | *ARHGEF7* | 25.2 | 1 | 0.188 | 0.0729 | 0.658 |
| HGNC:1034 | *BECN1* | 4.14 | 0.912 | 0.361 | 0.158 | 0.658 |
| HGNC:25180 | *FCHO2* | 19.7 | 0.999 | 0.234 | 0.102 | 0.668 |
| HGNC:9785 | *RAB5C* | 10.11 | 0.933 | 0.367 | 0.0774 | 0.668 |
| HGNC:1554 | *CBX4* | 59.47 | 0.983 | 0.282 | 0.0595 | 0.671 |
| HGNC:29221 | *RALGAPB* | 12.23 | 1 | 0.106 | 0.0411 | 0.672 |
| HGNC:16397 | *OSBPL11* | 46.4 | 0.942 | 0.335 | 0.17 | 0.673 |
| HGNC:25642 | *RSBN1* | 8.91 | 0.131 | 0.435 | 0.249 | 0.688 |
| HGNC:25788 | *ARMH3* | 7.54 | 0.00000708 | 0.570 | 0.38 | 0.695 |
| HGNC:2170 | *CNTFR* | 12.62 | 0.954 | 0.342 | 0.109 | 0.699 |
| HGNC:9229 | *PLPP3* | 29.28 | 0.921 | 0.381 | 0.0804 | 0.704 |
| HGNC:6321 | *KIF3C* | 22.27 | 0.986 | 0.301 | 0.131 | 0.708 |
| HGNC:6551 | *LEF1* | 0.93 | 0.998 | 0.205 | 0.0433 | 0.71 |
| HGNC:12733 | *WASF2* | 12.98 | 0.963 | 0.332 | 0.105 | 0.712 |
| HGNC:17060 | *KIFAP3* | 8.31 | 0.0000456 | 0.536 | 0.353 | 0.719 |
| HGNC:8587 | *PAICS* | 6.17 | 0.0000244 | 0.842 | 0.509 | 0.719 |
| HGNC:23817 | *CNOT10* | 29.82 | 0.983 | 0.305 | 0.162 | 0.743 |
| HGNC:17851 | *REXO2* | 4.85 | 0.0836 | 0.719 | 0.314 | 0.754 |
| HGNC:18287 | *VPS37D* | 64.11 | 0.923 | 0.368 | 0 | 0.755 |
| HGNC:29091 | *ATG13* | 17.94 | 0.961 | 0.331 | 0.145 | 0.757 |
| HGNC:16703 | *SLC17A6* | 5.76 | 0.168 | 0.469 | 0.25 | 0.763 |
| HGNC:24523 | *ZZZ3* | 5.69 | 1 | 0.178 | 0.069 | 0.763 |
| HGNC:22229 | *ZMIZ2* | 50.33 | 1 | 0.183 | 0.071 | 0.77 |
| HGNC:29025 | *ZNF536* | 1.66 | 0.999 | 0.222 | 0.086 | 0.775 |
| HGNC:24670 | *FNDC3B* | 19.52 | 1 | 0.043 | 0 | 0.787 |
| HGNC:25945 | *ARL15* | 5.72 | 0.000934 | 1.210 | 0.611 | 0.79 |
| HGNC:3031 | *ARID3A* | 66.1 | 0.991 | 0.274 | 0.0872 | 0.794 |
| HGNC:30257 | *PYGO2* | 31.65 | 0.953 | 0.314 | 0 | 0.795 |
| HGNC:23713 | *ZNF496* | 81.67 | 0.996 | 0.249 | 0.0792 | 0.806 |
| HGNC:16971 | *FRS2* | 1.16 | 0.998 | 0.213 | 0.0449 | 0.807 |
| HGNC:6529 | *LCP2* | 44.22 | 0.979 | 0.311 | 0.148 | 0.824 |
| HGNC:24877 | *YIPF5* | 19.06 | 0.987 | 0.229 | 0 | 0.829 |
| HGNC:18807 | *CAMTA2* | 36.88 | 0.998 | 0.271 | 0.16 | 0.846 |
| HGNC:6021 | *IL6ST* | 41.53 | 1 | 0.197 | 0.0861 | 0.853 |
| HGNC:15720 | *STRN3* | 7.73 | 0.817 | 0.354 | 0.196 | 0.857 |
| HGNC:8971 | *PIK3C2A* | 6.4 | 0.00383 | 0.370 | 0.261 | 0.861 |
| HGNC:12569 | *UNC5C* | 8.83 | 0.000644 | 0.502 | 0.321 | 0.867 |
| HGNC:6040 | *ILK* | 2.82 | 0.0000182 | 0.771 | 0.476 | 0.898 |
| HGNC:18222 | *RBFOX1* | 0.34 | 0.955 | 0.335 | 0.146 | 0.96 |

| | | | | | | |
|---|---|---|---|---|---|---|
| HGNC:29858 | *NLK* | 5.09 | NA | NA | NA | NA |

#### Supplementary Table 8. Clinical description of patients with variants in *VPS4.*

| | 100KGP patient 1 | 100KGP patient 2 | CMG patient |
|---|---|---|---|
| *de novo* variant | 16:69320768:A:T (GRCh38) | 16:69319539:G:A (GRCh38) | Variant data unavailable |
| Behavioural phenotypes | • **Intellectual disability, profound**<br>• **Profound global developmental delay**<br>• **Severe receptive language delay** | • **Intellectual disability**<br>• **Global developmental delay**<br>• **Delayed speech and language development**<br>• Abnormality of prenatal development or birth | • **Psychosocial retardation** |
| Movement / Muscle phenotypes | • **Delayed fine motor development**<br>• **Delayed gross motor development**<br>• Generalized hypotonia<br>• Generalized dystonia<br>• Chorea | • **Delayed fine motor development**<br>• **Delayed gross motor development**<br>• Inability to walk | • Right spastic hemiparesis |
| Seizure phenotypes | | • **Seizures** | • **Epilepsy** |
| Other brain phenotypes | • **Congenital microcephaly**<br>• Frontoparietal polymicrogyria | • **Microcephaly**<br>• Morphological abnormality of the central nervous system | • **Microcephaly**<br>• Frontoencephalocele |
| Other phenotypes | • **Poor visual behavior for age**<br>• Esophagitis | • **Abnormality of the eye**<br>• **Developmental cataract**<br>• Talipes<br>• Abnormality of male external genitalia | |

#### Supplementary Table 9. Clinical description of patients with *de novo* variants in *TMEM63B*.

| | DDD patient 1 | 100KGP patient 1 | 100KGP patient 2 | 100KGP patient 3 | 100KGP patient 4 |
|---|---|---|---|---|---|
| *de novo* variant | 6:44134714:G:A (GRCh38) | | | 6:44151868:G:A (GRCh38) | 6:44148860:TCC:: (GRCh38) |
| Behavioural phenotypes | • Abnormality of the nervous system | • **Intellectual disability**<br>• **Global developmental delay**<br>• **Delayed speech and language development** | • **Intellectual disability, severe** | • **Mild global developmental delay**<br>• Hyperactivity | • **Intellectual disability, profound**<br>• **Global developmental delay**<br>• **Delayed speech and language development** |
| Movement phenotypes | • Abnormality of the nervous system | • **Delayed gross motor development**<br>• **Inability to walk**<br>• Delayed fine motor development | • Generalized hypotonia<br>• Abnormality of movement | • Clumsiness<br>• Falls | • **Delayed gross motor development**<br>• **Inability to walk** |
| Seizure phenotypes | • Abnormality of the nervous system | • **Seizures** | • **Focal-onset seizure**<br>• **Generalized-onset seizure**<br>• **Infantile spasms**<br>• **EEG with focal epileptiform discharges**<br>• **EEG with generalized epileptiform discharges**<br>• **EEG with burst suppression** | | • **Seizures** |
| Other brain phenotypes | • Abnormality of the nervous system<br>• Abnormality of head or neck | • Microcephaly<br>• Morphological abnormality of the central nervous system | • Infantile encephalopathy | • Cerebral hypomyelination<br>• Cerebral white matter hypoplasia<br>• Diffuse white matter abnormalities | • Progressive macrocephaly<br>• Severe hydrocephalus |
| Other phenotypes | • **Growth abnormality**<br>• Abnormality of the skeletal system<br>• Abnormality of abdomen morphology<br>• Abnormality of blood and blood-forming tissues<br>• Abnormality of metabolism/homeostasis<br>• Abnormality of the immune system<br>• Abnormality of the ear | • **Abnormality of the eye** | • Large for gestational age<br>• **Tall stature**<br>• Prominent eyelashes<br>• Broad eyebrow | • **Strabismus**<br>• Supernumerary nipple<br>• Cafe-au-lait spot<br>• Abnormal hair pattern | |

**Supplementary Table 10. FUSIL categories.**

| Mouse viability phenotype | Human cell essentiality score | FUSIL category | Class | Number of genes |
|---|---|---|---|---|
| Lethal | ≤ -0.45 | **Cellular lethal (CL)** | Lethal in mouse and essential in human cell lines | **413** |
| | > -0.45 | **Developmental lethal (DL)** | Lethal in mouse but non-essential in human cell lines | **764** |
| Subviable | > -0.45 | **Subviable (SV)** | Subviable in mouse and non-essential in human cell lines | **421** |
| | ≤ -0.45 | Subviable outlier (SV.outlier) | Subviable in mouse and essential in human cell lines | 16 |
| Viable | > -0.45 | **Viable with phenotype (VP)** | Viable and non-essential in human cells (at least one significant phenotype it in the adult homozygous null mice) | **1,867** |
| | > -0.45 | **Viable with no phenotype (VN)** | Viable and non-essential in human cells (no significant phenotype hits in the adult homozygous null mice when % procedures done ≥ 50%) | **318** |
| | > -0.45 | Viable insufficient data on procedures (V. insuffProcedures) | Viable and non-essential in human cells (no significant phenotype hits in the adult homozygous null mice when % procedures done < 50% / difficult to ascertain) | 625 |
| | ≤ -0.45 | Viable outlier (V.outlier) | Viable in the mouse & essential in human cells | 22 |

**Supplementary Table 11. IMPC and MGI viability assessment.**

| IMPC Viability (primary viability assessment) | MGI Viability (reported phenotypes) | Number of genes | % of discrepancy with respect to IMPC Viability category |
|---|---|---|---|
| Lethal | Lethal | 504 | |
| Lethal | Viable | 63 | 11.11% |
| Subviable | Lethal | 141 | - |
| Subviable | Viable | 110 | - |
| Viable | Lethal | 154 | 11.87% |
| Viable | Viable | 1,143 | |

#### Supplementary Figures

**Supplementary Fig. 1. Selection of mean Avana score threshold to identify essential genes. a) and b) Distribution of IMPC viability categories across bins of mean Avana scores.** Percentage (a) and numbers (b) of lethal, subviable and viable mouse-to-human orthologous genes across mean Avana score bins comprising 4,446 genes for which there was IMPC viability data, a good confidence orthologue and an Avana viability score (release 18Q3 of August 2018 for 17,634 genes in 485 cell lines). For genes with an Avana mean score below -0.45, the mouse null homozygotes were lethal in almost all cases, while genes with an Avana mean score above -0.45 presented lethal, subviable or viable phenotypes. **c) IMPC Viability categories across 11 cell lines.** A similar pattern was observed when a different source consisting on 11 cell lines from 3 different studies was used (threshold criteria explained in Munoz-Fuentes, et al. [19]. **d) F1 scores for the comparison with previous datasets.** F1 scores derived from the confusion matrices considering different Avana mean scores and the classification in essential versus non-essential genes from previous studies. An Avana score cut-off of -0.45, which maximises the F1 scores across the different datasets, was selected, so that all genes with an Avana mean score below or equal to -0.45 were considered essential.

**Supplementary Fig. 2. Distribution of different constraint scores derived from human population sequencing data across the five FUSIL categories established in this study. a), b) and c) Observed versus expected (o/e) ratio of gnomAD 2.1 scores. (a)** Distribution of o/e LoF scores; lower scores indicate more intolerance to LoF (outliers not shown). **(b)** Distribution of o/e missense scores; lower scores indicate more intolerance to missense variation (outliers not shown). **(c)** Distribution of o/e synonymous scores; lower scores indicate more intolerance to synonymous variation (outliers not shown). **d) Estimates of selection against heterozygous loss of gene function (shet).** The selective effects for heterozygous protein-truncating variants (shet) were obtained from the supplementary material of Cassa, et al. [12], with higher values indicating more intolerant to variation (outliers not shown). **d) Residual Variance Intolerance Score (RVIS).** Distribution of the Residual Variation Intolerance Score (RVIS; version CCDSr20)[5], with lower values indicating more intolerance (outliers not shown). **e) Haploinsufficiency percentage score (HI%).** Haploinsufficiency score as a percentage, computed by the Deciphering Developmental Disorders (DDD) consortium. High ranks (e.g. 0-10%) indicate a gene is more likely to exhibit haploinsufficiency, low ranks (e.g. 90-100%) indicate a gene is more likely to NOT exhibit haploinsufficiency. Significance of pairwise comparisons for all the constraint metrics are shown in Supplementary Table 5.

**Supplementary Fig. 3. HPO phenotypes for *VPS4A* cases.** The set of HPO encoded phenotypes reported for each case listed in Supplementary Table 8 was plotted as a subgraph of the ontology. Uninformative terms (those annotated to the same objects as all their children) were removed. a: 100KGP patient 1, b: 100KGP patient 2, c: CMG patient. The colour indicates whether the phenotype has been observed in 3 (orange), 2 (blue) or only 1 (grey) patient. For patient c, the original reported phenotypes were replaced by either the synonymous term or the closest term in the HPO (seizures: epilepsy; fontal encephalocele: frontoecephalocele; spastic hemiparesis: right spastic hemiparesis; delayed social development: psychosocial retardation).

**Supplementary Fig. 4. HPO phenotypes for *TMEM63B* cases.** The set of HPO encoded phenotypes reported for each case listed in Supplementary Table 9 was plotted as a subgraph of the ontology. Uninformative terms (those annotated to the same objects as all their children) were removed. a: DDD patient 1, b: 100KGP patient 1, c: 100KGP patient 2, d: 100KGP patient 3, e: 100KGP patient 4. The colour indicates whether the phenotype has been observed in 5 (dark orange), 4 (orange), 3 (dark blue), 2 (blue) or only 1 (grey) patient.

**Supplementary Fig. 5. Evidence for mouse viability for the genes considered in this study (a; Table 1, Methods) and genes annotated in MGI (b). a) IMPC Viability.** Percent distribution of primary viability assessment outcomes as obtained from the IMPC. **b) MGI**

**Viability.** Percent distribution of viability annotations as obtained from Mouse Genome Informatics (MGI)[56] for the same genes. Gene to phenotype annotations (excluding conditional annotations) from MGI were used to identify the set of genes with embryo lethality phenotypes (50 Mammalian Phenotype Ontology terms as described in Dickinson, et al. [23]; viability outcomes inferred from MGI annotations do not include the IMPC subviable category**. c) Correspondence between IMPC and MGI annotations.** For each IMPC viability category, the bar plots represent the percentage distribution of the viability assessment according to MGI annotations. For 2,115 mouse genes with both IMPC and non-IMPC phenotypic annotations available to infer viability, we found discrepancies for a set of 63 genes that were found to be lethal according to the IMPC but had no previous records of lethality in MGI as well as for 154 genes viable as reported by the IMPC and with some type of lethality annotations reported in MGI (10% overall discrepancy).

### Supplementary Fig. 1

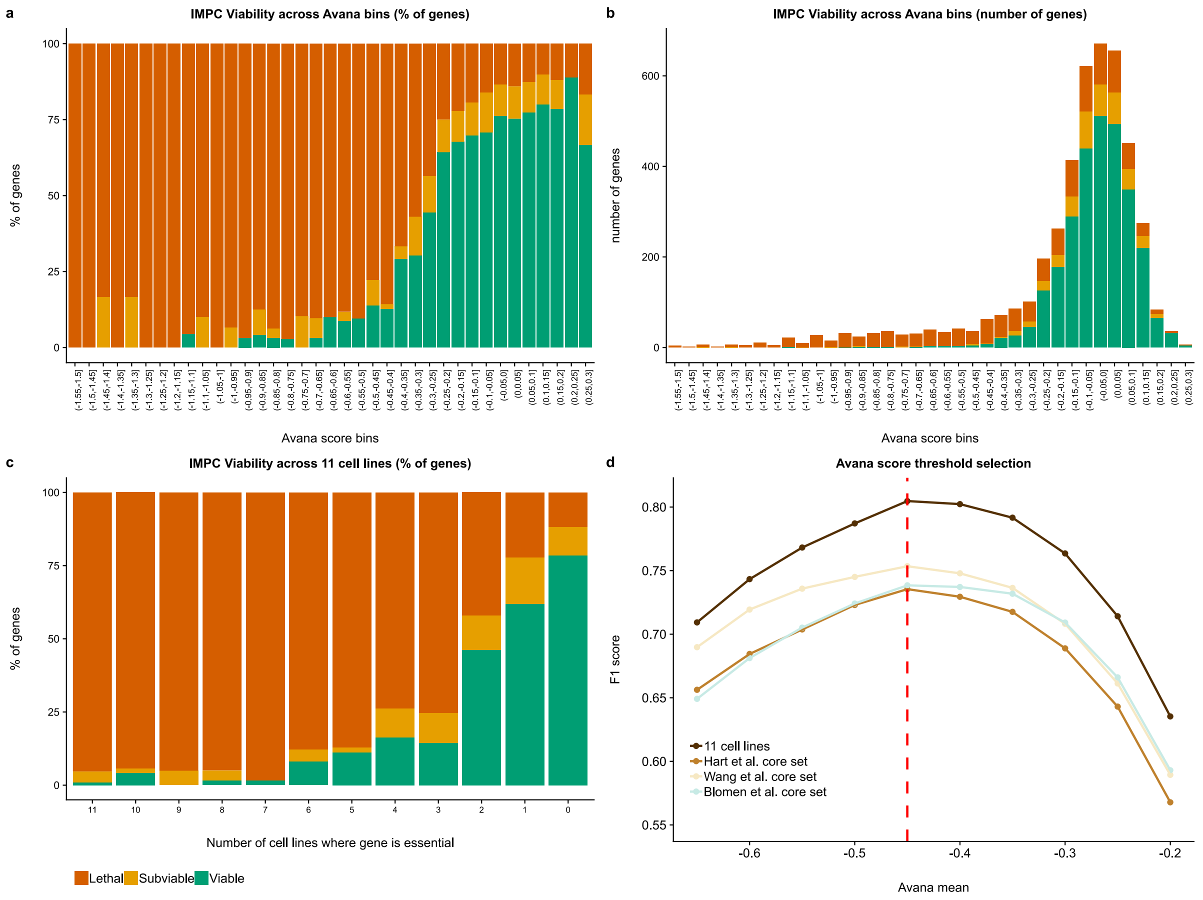

#### Supplementary Fig. 2

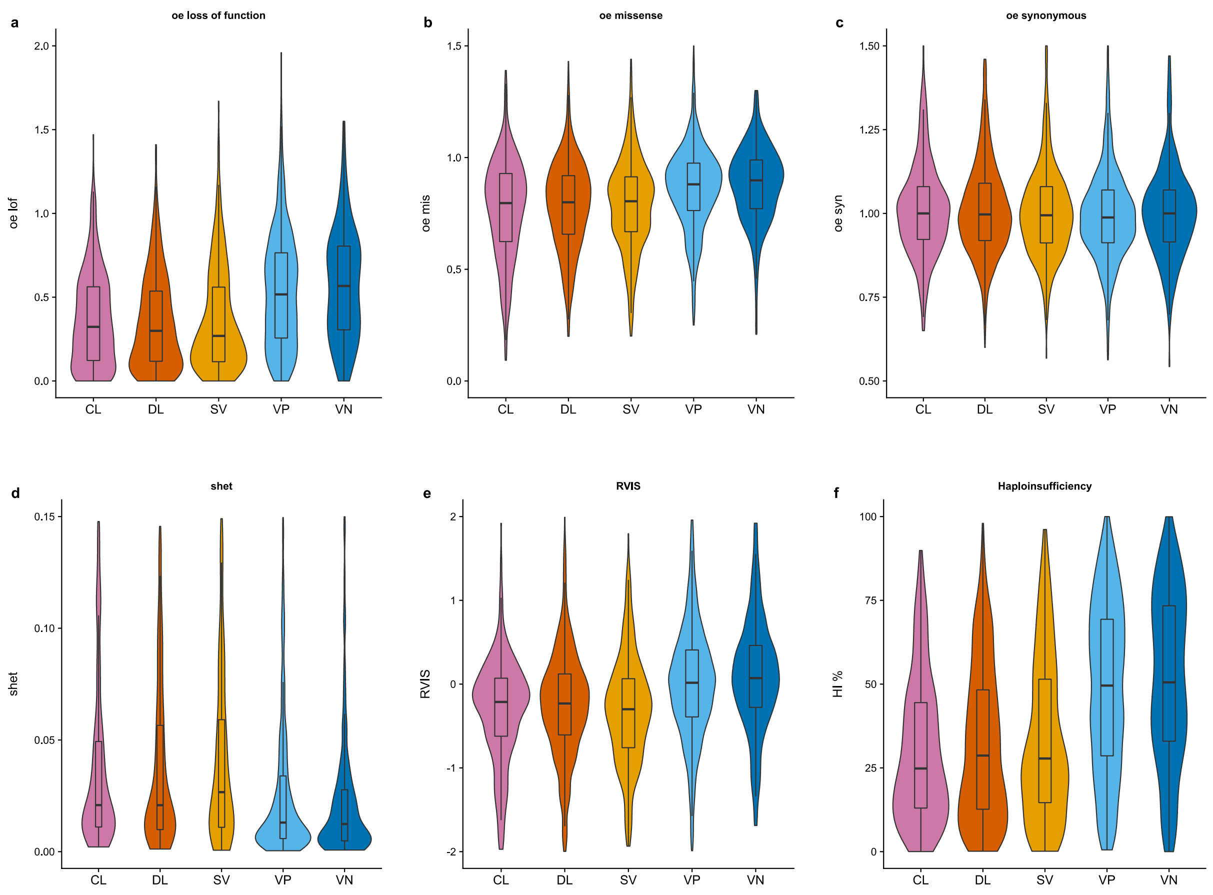

#### Supplementary Fig. 3

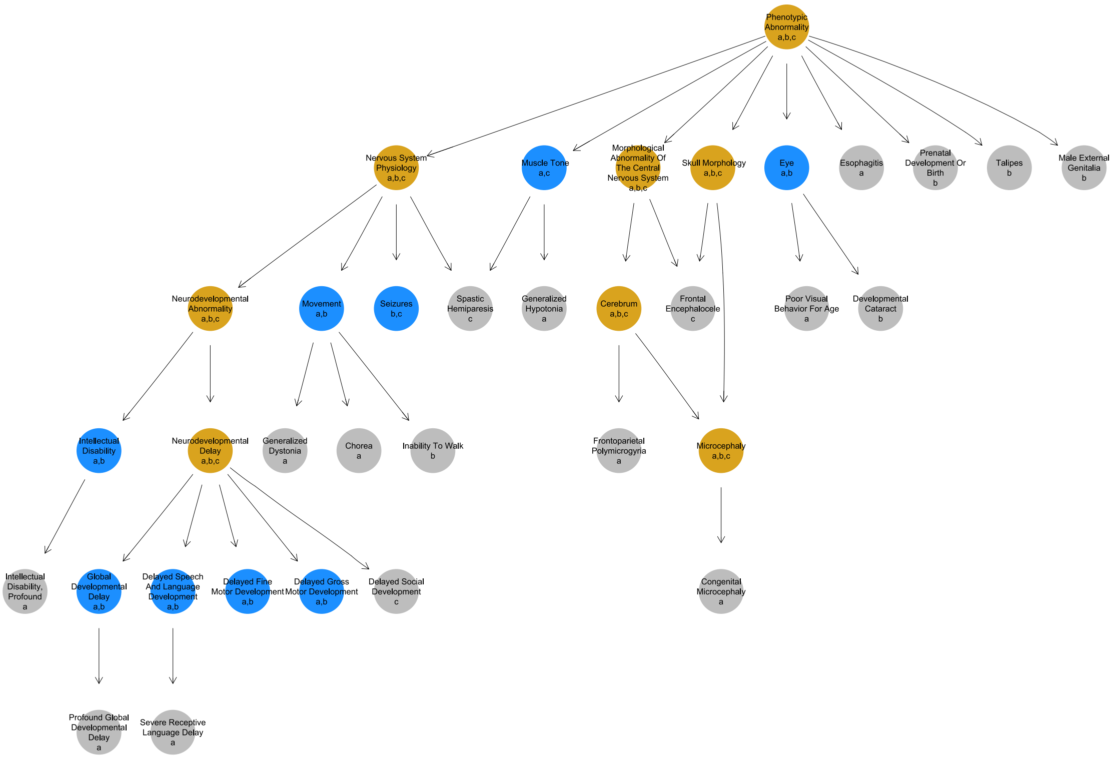

### Supplementary Fig. 4

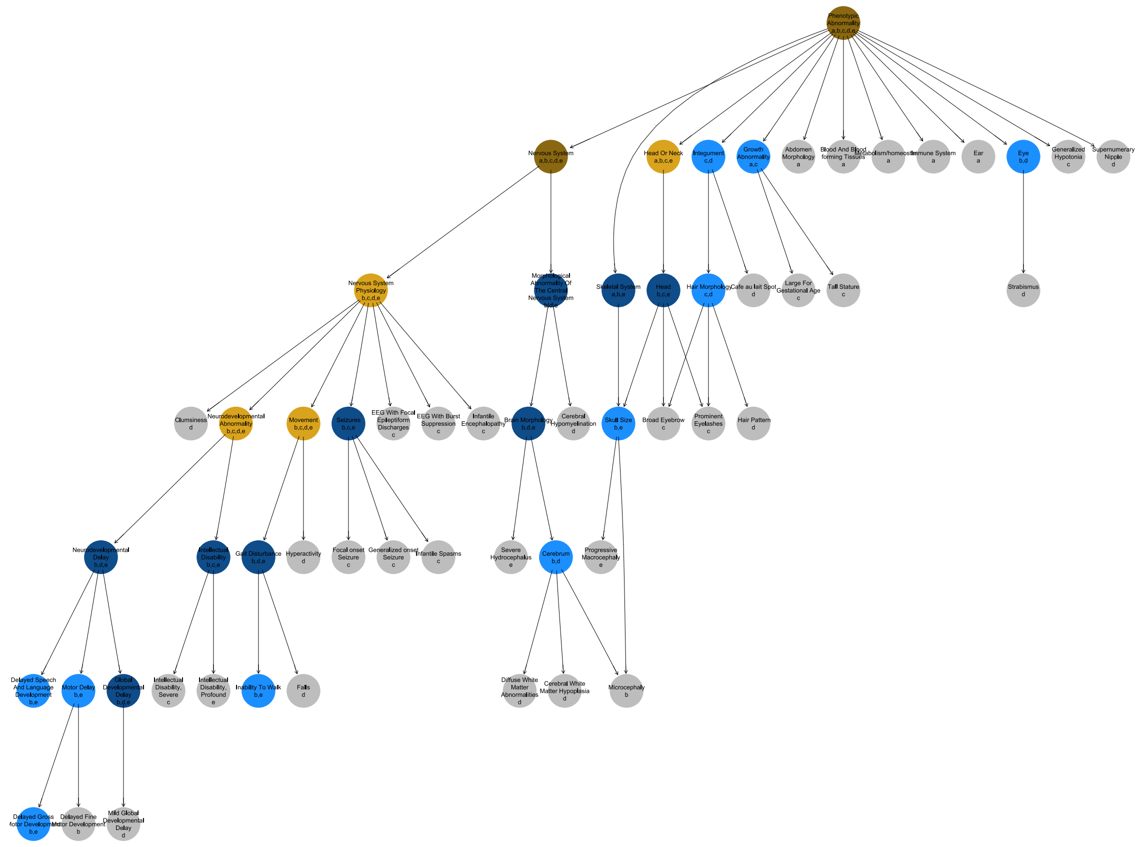

#### Supplementary Fig. 5

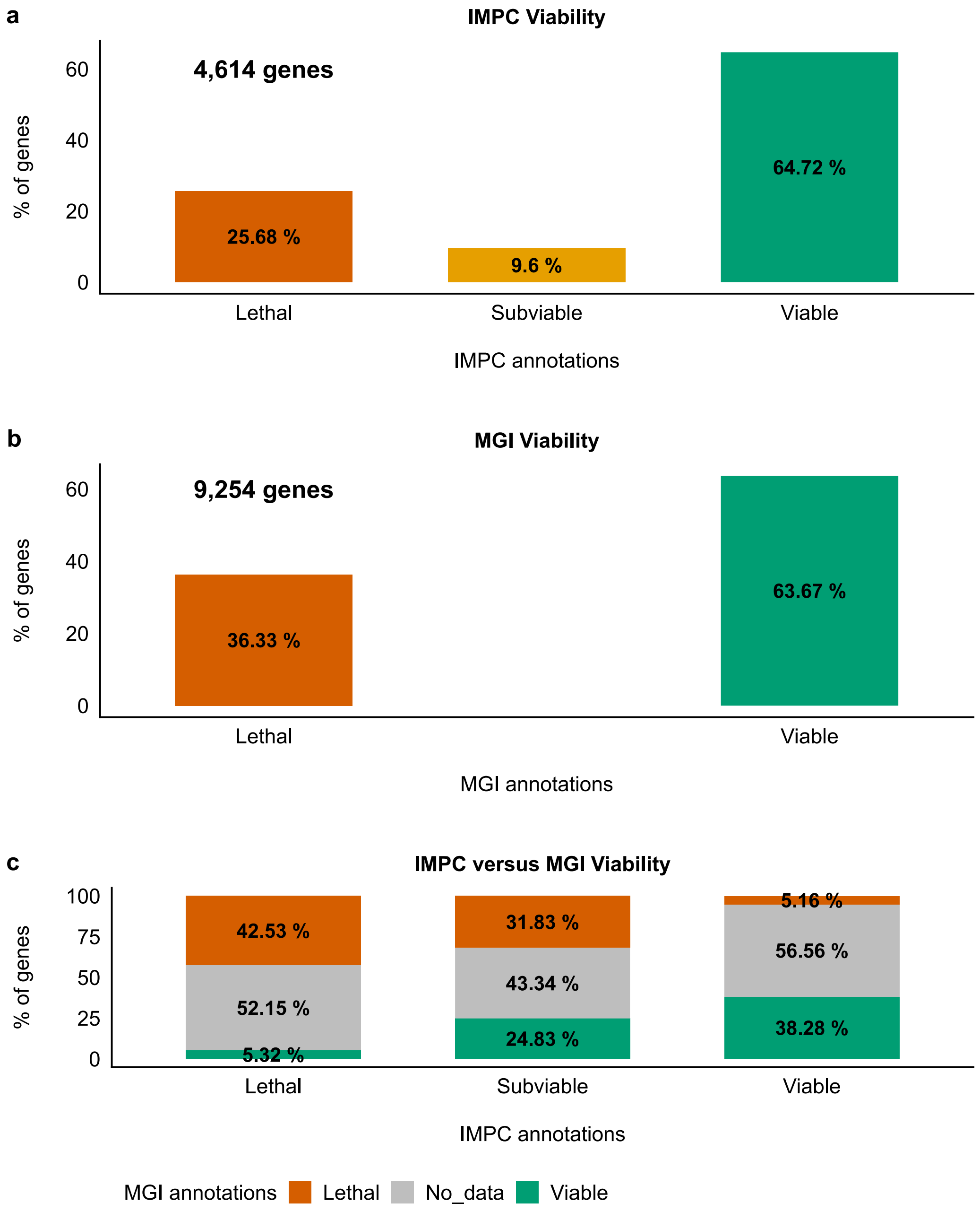
